# Precisely patterned nanofibers made from extendable protein multiplexes

**DOI:** 10.1101/2022.10.14.511843

**Authors:** Neville P. Bethel, Andrew J. Borst, Fabio Parmeggiani, Matthew J. Bick, TJ Brunette, Hannah Nguyen, Alex Kang, Asim K. Bera, Lauren Carter, Marcos C. Miranda, Ryan Kibler, Mila Lamb, Xinting Li, Banumathi Sankaran, David Baker

## Abstract

Molecular systems with coincident cyclic and superhelical symmetry axes have considerable advantages for materials design as they can be readily lengthened or shortened by changing the length of the constituent monomers. Among proteins, alpha helical coiled coils have such symmetric extendable architectures, but are limited by the relatively fixed geometry and flexibility of the helical protomers. Here, we describe a systematic approach to generating modular and rigid repeat protein oligomers with coincident C_2_ to C_8_ and superhelical symmetry axes that can be readily extended by repeat propagation. From these building blocks, we demonstrate that a wide range of unbounded fibers can be systematically designed by introducing hydrophilic surface patches that force staggering of the monomers; the geometry of such fibers can be precisely tuned by varying the number of repeat units in the monomer and the placement of the hydrophilic patches.

---

Both cyclic symmetry and superhelical symmetry are frequent in nature, but few systems have both cyclic and internal superhelical symmetry with coincident symmetry axes (Figure 1a). This geometry has the advantage that the individual protomer can be readily extended based on the internal superhelical symmetry such that the newly added portion makes the same interactions with its cyclic symmetric counterparts as the original protomer made with its counterparts. Amongst protein systems, coiled coils and the collagen triple helix have this very useful property, which has been widely exploited in natural biological systems and in protein engineering (1). However, the single stranded nature of these structures has limitations: the monomers are flexible and not readily amenable to protein fusion, the assemblies are restricted to a narrow range of twist and radius, and cannot readily be stacked along the axis of extension due steric constraints. The superhelical symmetry necessary for forming such structures is also found in helical repeat proteins, both natural and designed, composed of a globular protein unit that is tandemly repeated to form a rigid structure. *De novo* helical repeat (DHR) proteins have potential advantages as protomers over single helices, as they are rigid and amenable to protein fusion, can adopt a wide variety of geometries (2), and can stack using head-to-tail interactions similar to nucleic acids. However, while homo-oligomers have been generated using DHRs (3), the cyclic axes of the oligomer and the superhelical axes of the monomers have not been coincident, so extending the monomer does not extend the homo-oligomeric interface as is the case in coiled coils and double stranded nucleic acids.

**Figure 1:**
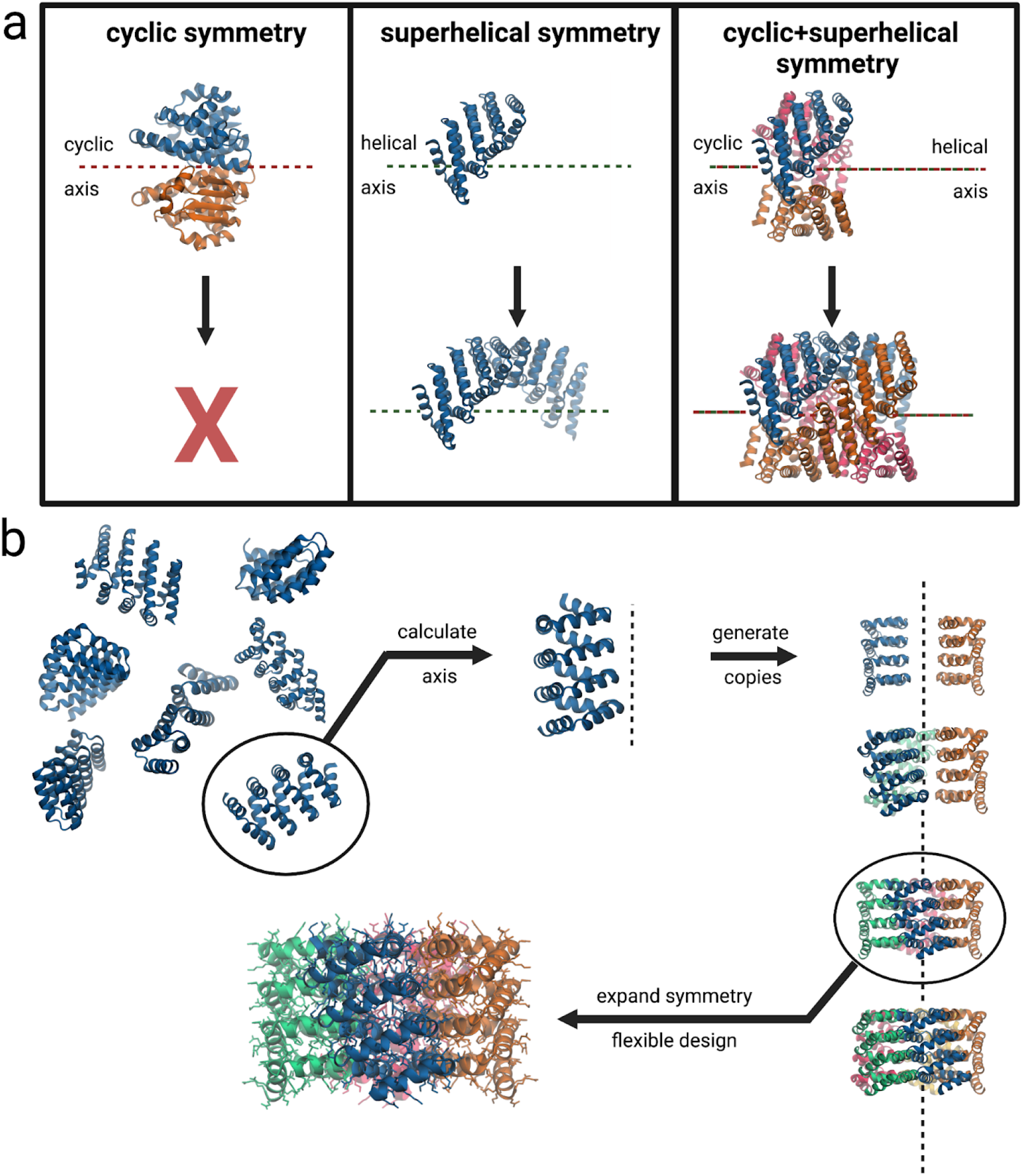
Concept and design approach. a. Examples of cyclic symmetry, superhelical symmetry, and the combination of both cyclic and superhelical symmetry. b. Multiplexes are created by first generating *de novo* repeat protein monomers through backbone fragment assembly. The monomer superhelical axis is calculated and evenly spaced copies are generated around the axis. Different cyclic symmetries are attempted, and specific symmetries are selected according to contact number and clash score. Filtered backbones are then assigned sequences before further computational filtering and experimental characterization.

We set out to systematically generate protein nanostructures with shared cyclic and superhelical symmetry axes based on cyclic helical repeat proteins (CHRs). We began by using fragment assembly to generate a wide variety of repeat protein monomers with repeat units with square two helix or triangular three helix geometry and four repeats in total (Figure 1b). The superhelix traced out by the centroids of the repeat units was computed, and from two to eight copies of the monomer were placed around the superhelical axis. Cyclic assemblies lacking backbone clashes and with extensive helix-helix intermolecular contacts were then computationally assigned sequences. We initially used Rosetta packing movers such as FastDesign and PackRotamers (4) but obtained better experimental success rates using proteinMPNN, a message passing neural net trained to predict sequences that will fold into a given protein structure (5). We selected subsets of designed multiplexes that had backbone configurations that closely matched predictions from either Alphafold2 or Alphafold multimer (Extended Data Figure 1) (6,7), and obtained synthetic genes for experimental characterization.

We expressed 67 of the proteinMPNN designed multiplexes in E coli, and characterized their oligomerization state by size exclusion chromatography (SEC). Of these designs, 60 were soluble and 11 of the 67 were monodisperse with elution profiles consistent with the oligomerization state, which was further confirmed by SEC-MALS measurements (Figure 2) (Extended Data Table 1). Small angle x-ray scattering (SAXS) profiles (Figure 2) were close to those computed from the computational design models (8,9,10). The volatility ratio (V_r_) for each pair of curves (Extended Data Table 2) (11) (a better determinator of goodness of fit than χ^2^ since it is less dominated by the fitting at the Guinier region) was less than 12.6 in the range of values determined for other previously designed protein oligomers (12). The C_4_ to C_8_ designs were also imaged by negative stain EM, which further confirmed that the particles are monodisperse with the correct shape and size (Extended Data Figure 2).

**Figure 2:**
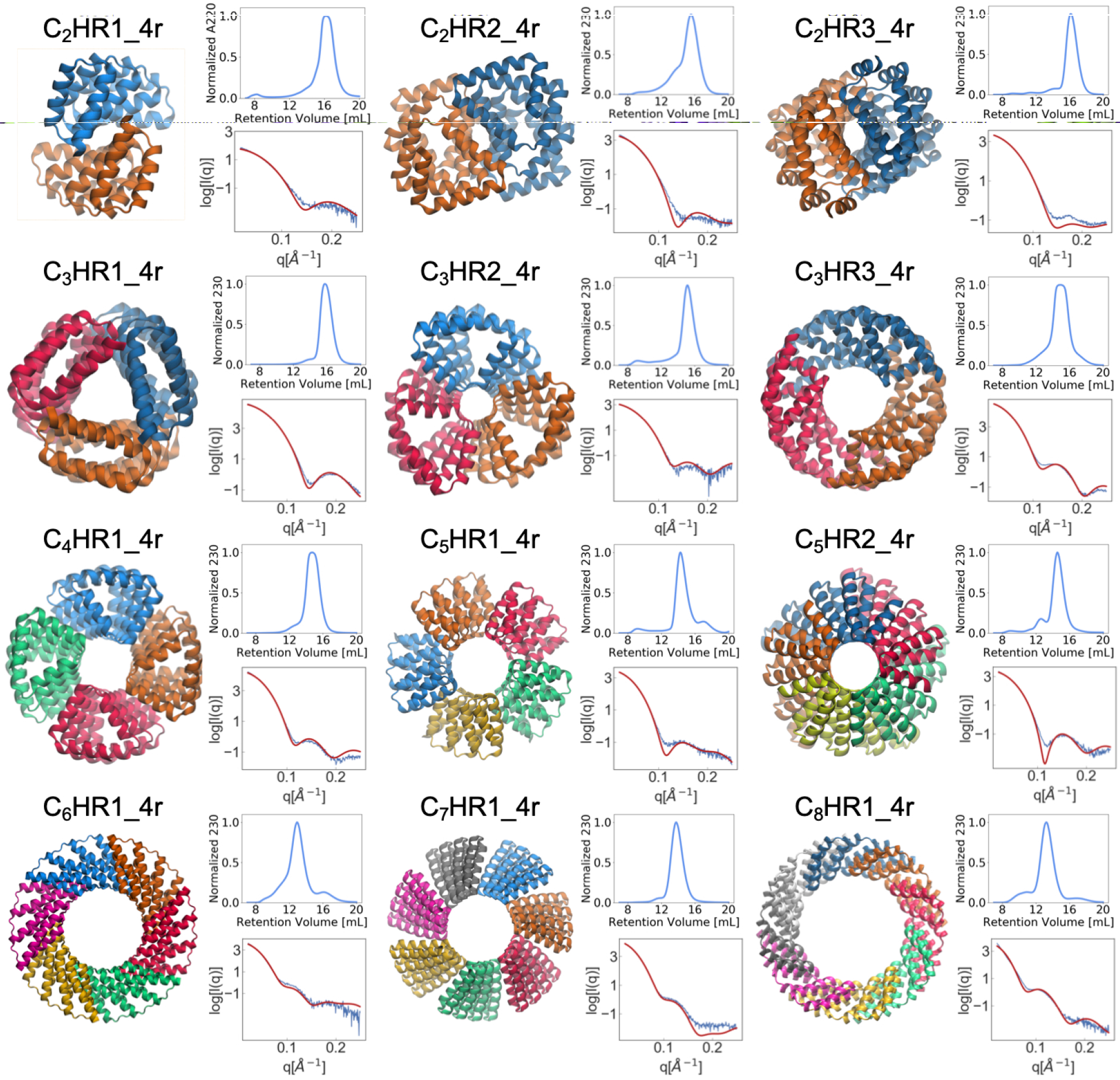
Experimentally validated multiplexes. The design models are each shown with their cyclic axes pointing into the page. For each design, the top right panel shows their size exclusion chromatography curve after IMAC purification and the bottom panel shows experimental (blue line) and model fit (red line) SAXS curves.

We determined the high resolution structures of five designs from C_2_ to C_6_ symmetry by x-ray crystallography, cryogenic electron microscopy (cryoEM) or both (Figure 3) (Extended Data Table 3). The crystal structure of C_2_HR1_4r matches the design model with an overall backbone RMSD of 1.55 Å. C_2_HR1_4r has a large repetitive interface composed primarily of leucines (Extended Data Figure 3a). For C_3_HR3_4r, the overall backbone RMSD between crystal structure and design model is 1.25 Å. In contrast to C_2_HR1_4r, C_3_HR3_4r has a large twist, causing the interface to become staggered with the top two repeats packing on the bottom two repeats of the adjacent monomer (Extended Data Figure 3b). C_4_HR1_4r was solved at 3.1 Å by x-ray crystallography and ∼3.7 Å by cryoEM; the two experimental structures are very close to the design model and to each other (RMSDs of 1.47 Å and 1.32 Å, respectively) with triangular shaped repeat monomers with inner cavities lined by phenylalanines (Extended Data Figure 3c). The C_5_HR2_4r interface is focused near the inner radius of the structure, and is composed of a thin strip of hydrophobic residues along the helical axis; desolvated salt bridges also line the inner radius of C_5_HR2_4r (Extended Data Figure 3d). C_5_HR2_4r has the largest rise of the structures, with an average rise of 1.1 nm per repeat. C_5_HR2_4r matches the design model with an overall backbone RMSD of 2.15 Å and has a V_r_ of 12.6, higher than that of the other designs, further suggesting that all 12 designs are close to the correct structure. C_6_HR1_4r matches the design model with an overall RMSD of 1.95 Å. Like C_5_HR2_4r and many of the other two helix repeat oligomers, the monomers of C_6_HR1_4r interact at the inner radius of the oligomer, but the repeats fan out towards the outer radius. The C_6_HR1_4r interface may be stabilized by a repetitive cation-pi interaction between tyrosine and arginine side chains of adjacent monomers (Extended Data Figure 3e). The high resolution structure of C_6_HR1_4r is the widest with an outer radius of 92 Å. The inner radius is 41 Å, which is large enough to fit a C_2_HR dimer.

**Figure 3:**
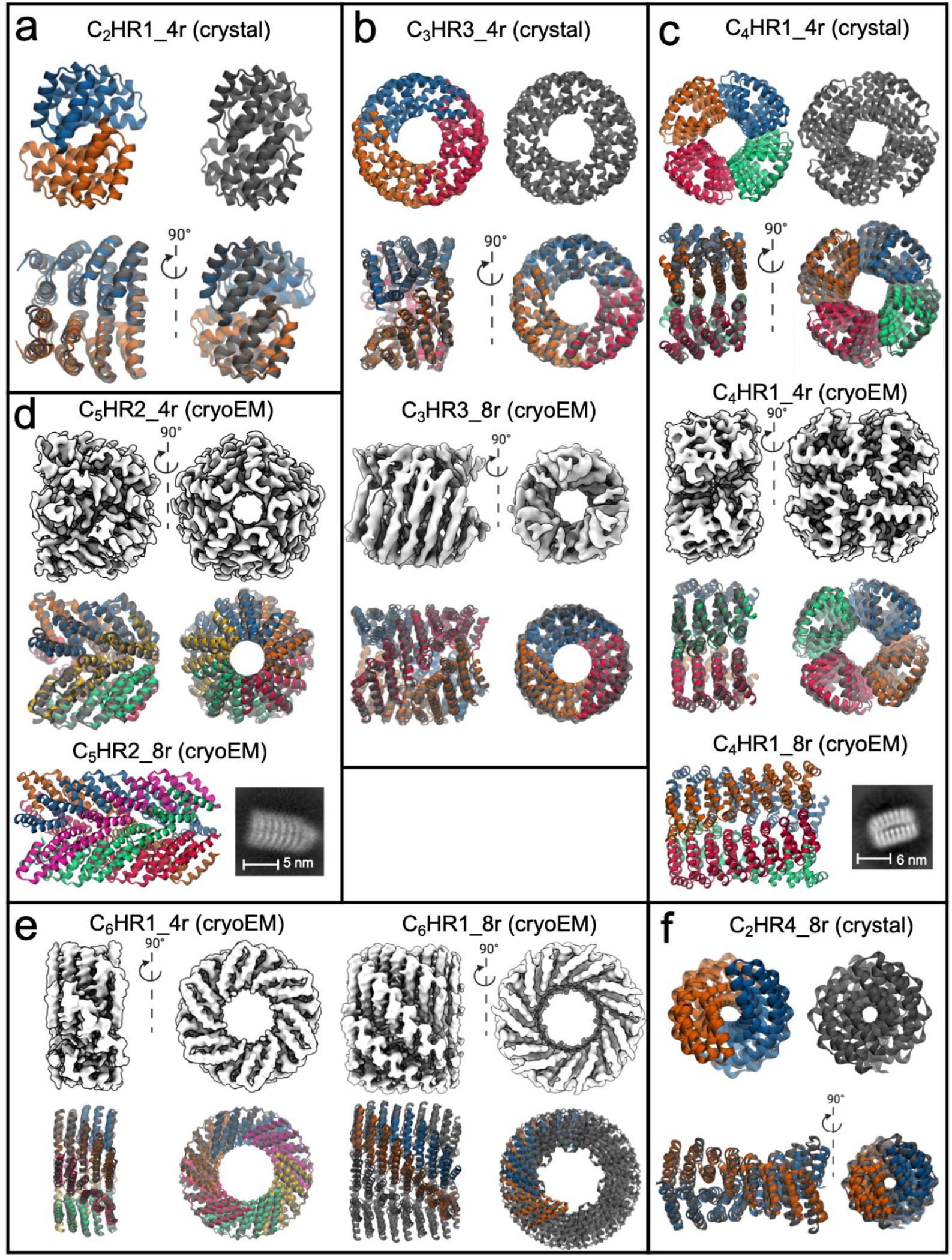
a. Crystal structure of C_2_HR1_4r aligned to design models by backbone RMSD. b. Crystal structure of C_3_HR3_4r and cryoEM model of C_3_HR3_8r. c. Crystal (upper) and cryoEM (middle) structures of C_4_HR1_4r. At the bottom the C_4_HR1_8r design model is shown as a side view with a corresponding cryoEM class average shown on the right. d. CryoEM structure of C_5_HR2_4r. At the bottom C_5_HR2_8r is shown as a side view with a corresponding cryoEM class average shown on the right. e. CryoEM structures of C_6_HR1_4r and C_6_HR1_8r. f. Crystal structure of C_2_HR4_8r. For all displayed structures, the experimentally determined structures are shown in gray while the backbone aligned design models are colored by chain. Since the C_6_HR1_8r cryoEM model is C_7_ instead of C_6_, only two chains of the design model were superimposed.

## Designed CHR multiplexes are extendable

Extendability is in principle a major advantage of helical repeat protein oligomers that have aligned superhelical and cyclic symmetry axes. Like the DNA duplex, they can geometrically be extended by propagating the number of repeats, and the interfacial contacts should increase with each additional repeat (Figure 1a). To investigate such extendability, we designed eight repeat versions of four of the validated four repeat multiplexes. The backbones of these proteins were propagated parametrically, and the sequences were designed similarly to the original four repeat versions. The SEC purified proteins form monodisperse particles with the expected size by SEC-MALS and expected shape as confirmed by negative stain electron microscopy (nsEM) and cryoEM (Figure 3b-f). The structures of the C_3_HR3_8r and C_6_HR1_8r were further analyzed by three dimensional reconstruction from cryoEM. Like C_3_HR3_4r, C_3_HR3_8r closely matches the design model with an overall backbone RMSD of 2.05 Å. SEC-MALS indicates a mixture of C_6_/C_7_ oligomers for C_6_HR1_8r with majority C_6_; while both states are apparent in cryoEM, we were only able to successfully reconstruct the C_7_ state, which is the largest of all the monodisperse designs with a total size of 305 kDa. The RMSD of the single monomer is 1.43 Å and two adjacent monomers is 2.74 Å, indicating that only subtle shifts in rotation and translation at the interface were required to accommodate the extra monomer. We also expressed and validated a C_2_ dimer directly as an eight repeat duplex (C_2_HR4_8r) that was verified by x-ray crystallography (RMSD to design model = 3.88 Å; Figure 3f). The twist of the crystal structure is approximately 22.5° per repeat or 16 repeats per turn, and hence by propagating this structure from 9 to 17 repeats, the angle between the N and C terminal repeats can be modulated from 180° to 360°.

## Design of patterned fibers by surface redesign

In principle, the monomers of the presented multiplexes could be extended indefinitely, but this is not currently feasible due to limitations on gene synthesis (increasing monomer length could also increase unintended higher order oligomers since the individual monomers can shift farther and farther along the helical axis). The top and bottom surfaces of the CHR monomers are primarily hydrophilic residues in the bounded multiplexes presented above. We attempted to replace these with the hydrophobic residues found on the core repeats but most of these “uncapped” designs did not express in *E.Coli*, likely due to the increased hydrophobicity and lack of specificity in forming the intended fiber geometry.

We instead adopted a “Lincoln logs” approach that alternates nonpolar patches that favor close subunit-subunit interactions with charged polar patches which disfavors buried interactions to introduce ‘pores’ on the fiber surface (Figure 4a). For the C_3_ to C_6_ designs, the monomers have four interfaces, which are composed of hydrophobic patches on the four corners of each monomer. The surface alternates from hydrophobic to hydrophilic to hydrophobic, which reduces nonspecific interfacial shifting and increases overall solubility.

**Figure 4:**
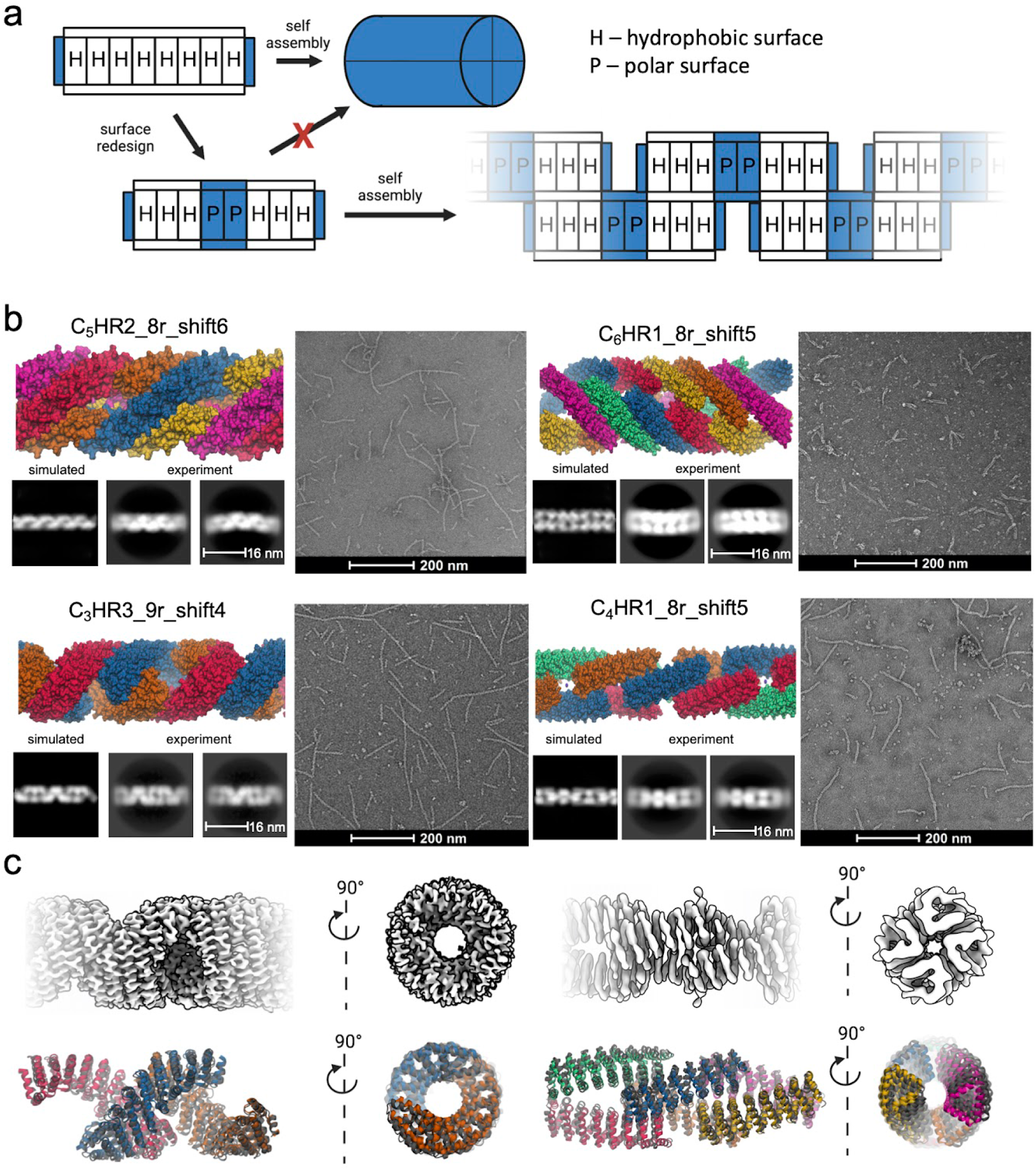
Patterned fibers. a. Concept for self assembling, patterned fibers. A single CHR monomer has two sides with hydrophobic surfaces. Dividing each surface with a hydrophilic patch forces a staggered formation, triggering fiber assembly. b. Examples of successfully assembled patterned fibers. Design models are shown on the upper left, negative stain micrographs on the right, and two dimensional class averages from negative stain are shown on the lower left. c. Three dimensional reconstructions from cryoEM data for C_3_HR3_9r_shift4 (left) and C_4_HR1_8r_shift5 (right). The upper row shows cryoEM densities of the symmetry expanded fibers. The lower row shows the design models aligned to the cryoEM structures. The cryoEM models are colored gray while the design models are colored by chain. The design models were aligned by backbone RMSD of the middle, blue chain.

We expressed and characterized 58 fibers redesigned from the four CHRs verified to be extendable. A total of 25 of the tested fibers assembled into 50 to 800 nm fibers readily observable by nsEM. The fibers are soluble, as we purified them through conventional IMAC and screened the elution fractions. The fibers grow over time (Extended Data Figure 4): for example C_3_HR3_9r_shift4 increases from approximately 100 nm to nearly a micron after three months incubation at 4°C; there are evidently kinetic traps during fiber assembly that can be overcome with sufficient time. We selected one fiber from each of the four CHRs for further characterization by nsEM (Figure 4b). The diameters of the fibers are closely consistent with the computational models, and the designed surface patterning closely matches the two dimensional class averages derived from nsEM.

We characterized the three dimensional structure of two of the fibers by cryoEM (Figure 4c). As with the bounded designs, the cryoEM model backbones closely match the design models. C_4_HR1_8r_shift5 has overall C2 symmetry; the fiber structure resembles chain links with each corresponding to a C_2_ unit. The pore size is approximately 880 Å^2^ (2 repeats). The fiber has a twist of 42.6° per monomer, or nearly 90° for every two monomers, and this feature along with the C_2_ symmetry can be exploited to generate square lattices of fibers. C_3_HR3_9r_shift4 was solved at 3.8 Å resolution, permitting more precise helix and sidechain assignment. The two interfaces of C_3_HR3_9r_shift4 are about the same size with approximately 38 carbon-carbon contacts (or 3 contacting repeats) per interface. The pore size of C_3_HR3_9r_shift4 is slightly larger (∼1020 Å^2^). Like C_4_HR1_8r_shift5, the pore size and twist closely match the design; the twist is -142.8° per monomer, or 71.4° per repeat. Notably, C_3_HR3 was verified to high resolution for the bounded four repeat, bounded eight repeat and unbounded fiber designs.

For materials engineering, a very useful aspect of our fiber design strategy is that the properties of the fibers can be tuned simply by changing the number of repeats units on the monomeric subunits, and the size of the hydrophilic spacer between the hydrophobic units forming the interface: the more hydrophobic units, the larger the subunit-subunit interface between monomers, and the larger the hydrophilic spacer, the larger the pores in the resulting fibers (Figure 5a). We explored varying both properties, and found that it is possible to lengthen the monomer one helix at a time, enabling control of the pore size of the fiber with single helix precision. 2D class averages indicate pore size and spacing consistent with the design models, with C_4_HR1_11r_shift7 having the largest pore size. We calculated the persistence lengths of the fibers from the nsEM data (Figure 5b) using the springEM software suite (13). Most of the fibers have persistence lengths around 2 microns, which is between the persistence lengths of intermediate filaments (500 nm) and actin (17.7 μm). The stiffest fiber by far is C_3_HR3_9r_shift4 with a persistence length of 7.44 microns. While the radii of all fibers are comparable, C_3_HR3 has the largest repeat size; thus, the mechanical stiffness of the monomer may be responsible for the increase in stiffness. For C_4_HR1, we expected that the stiffness would decrease with increasing pore size. Across the four C_4_HR1 variations, we find that this is indeed the case, with C_3_HR3_11r_shift7 having the lowest persistence length as measured by the springEM and as visualized by nsEM 2D class averaging of these assemblies.

**Figure 5:**
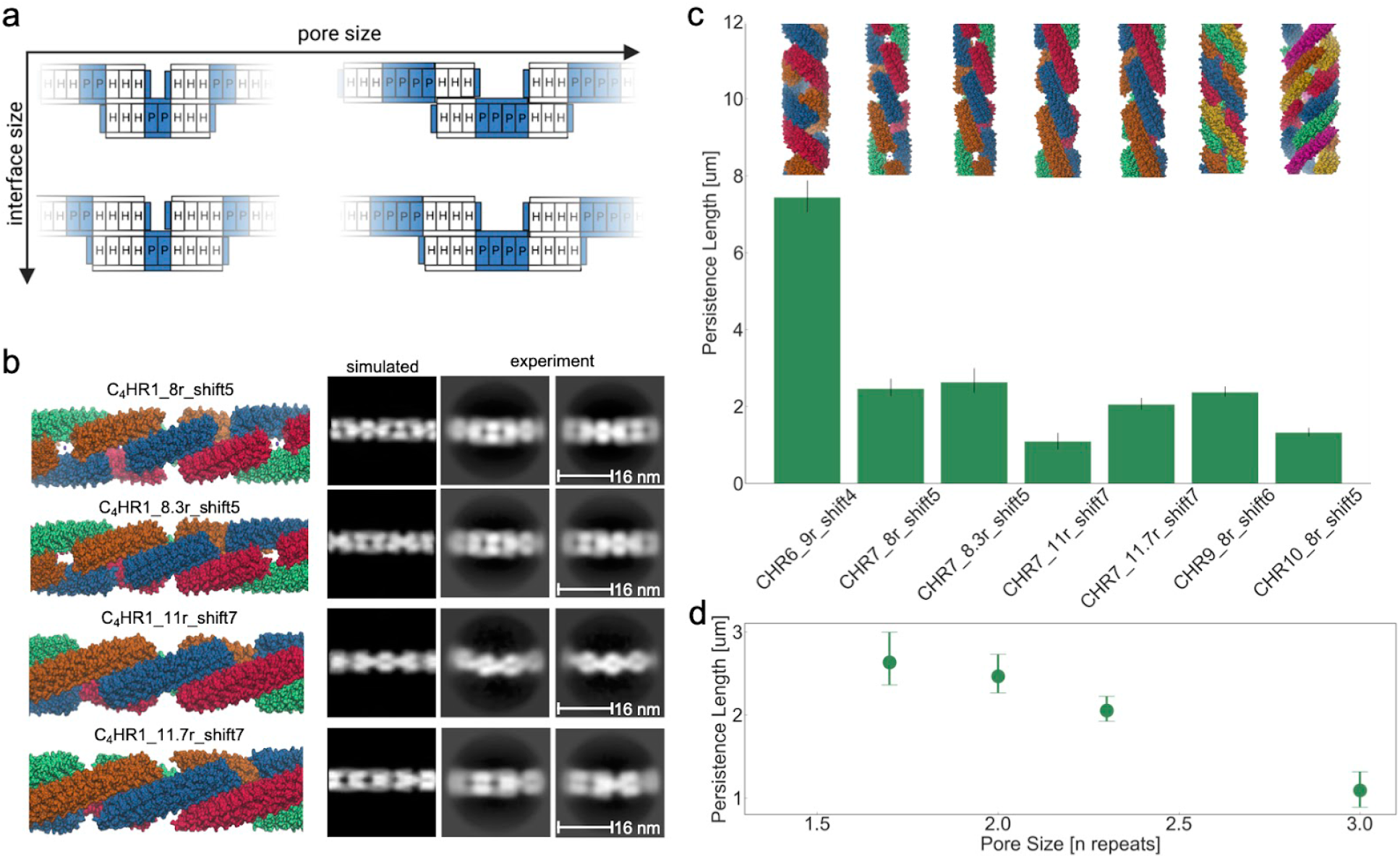
Sculpting the structures and persistence length of the patterned fibers. a. Concept for generating variations of patterned fibers. By changing the monomer size and spacing, fiber with different pore and interface size can be created. b. Variations of C_4_HR1 fibers. Chain length and spacing are varied between the fibers. Negative stain class averages are shown on the right. c. Persistence lengths of fibers calculated from negative stain micrographs. Error bars are 90% confidence intervals determined by bootstrap monte carlo. d. Persistence length over pore size for C_4_HR1 fibers.

## Discussion

Our designed assemblies with coincident cyclic and superhelical symmetry axes open up new frontiers in protein nanomaterial design. The designs span a wide range of monomer configurations, and are readily extendable by repeat propagation. By alternating the nonpolar monomer-monomer interaction regions with charged/polar surfaces having very large solvation free energy penalties for burial, protein filaments with different porosity and geometry can be robustly generated. The resulting porous structures could provide platforms for biomineralization analogous to collagen. The pores can also serve as binding sites for ligands containing one or two repeat units, enabling decoration of the fibers with molecules fused to these ligands at a readily tunable spacing. While here we primarily explore the assembly of one dimensional protein fibers, it should be possible to extend our approach to two to three dimensional materials. For example, the filaments could be resurfaced to form three dimensional lattices, or the bounded rings could be stacked in two dimensions to form extendable sheets. We show that the mechanical properties of the fibers can be modulated by changing the pore size of the fibers. Smaller pore sizes result in stiffer fibers, and this mechanism can be used to tune the mechanical properties of higher order materials built from the fibers.

## Acknowledgements

We thank Tim Huddy, Yang Hsia, Harley Pyles, Hao Shen, Alexis Courbet, Florian Praetorius and Lance Stewart for helpful discussions. We also thank Sasha Dickinson and Joel Quispe for training and operation of the electron microscopes at the Arnold & Mabel Beckman Center for Cryo-Electron Microscopy. We also thank Analisa Murray, Piper Heine, and Stacey Gerben for their work in expressing and purifying protein for crystal screens. We also thank the SYBYLIS group for their work on the SAXS data collection at the Advanced Light Source (ALS), a national user facility operated by Lawrence Berkeley National Laboratory on behalf of the Department of Energy, Office of Basic Energy Sciences, and Integrated Diffraction Analysis Technologies (IDAT) program. The SYBYLIS group is supported by the National Institute of Health project ALS-ENABLE (P30 GM124169) and a High-End Instrumentation Grant (S10OD018483). This work was supported with funds provided by the Howard Hughes Medical Institute (D.B.) and a Hanna Gray Postdoctoral fellowship (GT11817, N.P.B.), the Institute for Protein Design Directors Fund (MB), the Donald and Jo Anne Petersen Endowment for Accelerating Advancements in Alzheimer’s Disease Research (TJ.B.), and the Audacious Project at the Institute for Protein (A.J.B., H.N., A.K., M.C.M., L.C., M.L., X.L., R.K., D.B.). F.P. is the recipient of an EPSRC early career fellowship (EP/S017542/1) and was supported by the BrisSynBio grant BB/L01386X/1. This work was also supported, in whole or in part, by the Bill & Melinda Gates Foundation Grant # INV-010680. Crystallographic data was collected at the Advanced Light Source (ALS) and Advanced Photon Source (APS). Advanced Light Source (ALS) a national user facility operated by Lawrence Berkeley National Laboratory on behalf of the Department of Energy, Office of Basic Energy Sciences, through the Integrated Diffraction Analysis Technologies (IDAT) program, supported by DOE Office of Biological and Environmental Research. Additional support comes from the National Institute of Health project ALS-ENABLE (P30 GM124169) and a High-End Instrumentation Grant S10OD018483. At APS, the Northeastern Collaborative Access Team beamlines, which are funded by the National Institute of General Medical Sciences from the National Institutes of Health (P30 GM124165). This research used resources of the Advanced Photon Source, a U.S. Department of Energy (DOE) Office of Science User Facility operated for the DOE Office of Science by Argonne National Laboratory under Contract No. DE-AC02-06CH11357.

## Author Contributions

Conceptualization: F.P., N.P.B., D.B. Methodology: N.P.B. D.B. Software: N.P.B., A.J.B. T.B., R.K. Validation: N.P.B., A.J.B., H.N., A.K., A.K.B., M.C.M., L.C., M.J.B, M.L., X.L. Formal Analysis: N.P.B. A.J.B, A.K.B., M.J.B. Visualization: N.P.B., A.J.B. Supervision: A.J.B., D.B. Writing: N.P.B. D.B. Project Administration: N.B. Funding Acquisition: N.P.B, D.B

---

**Extended Data Figure 1:**
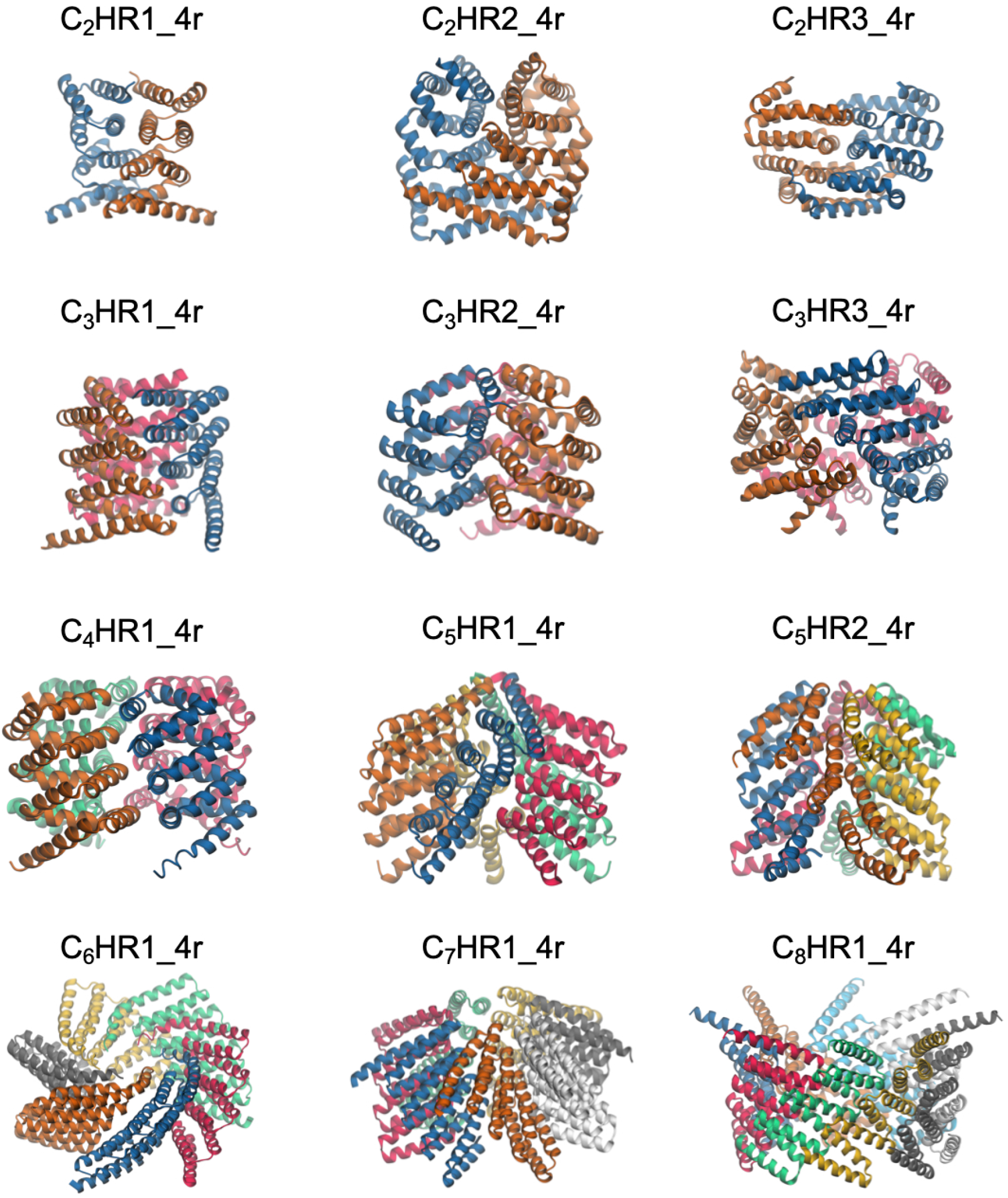
Alphafold2 models with histidine tags. These models were used in the SAXS analysis

**Extended Data Figure 2:**
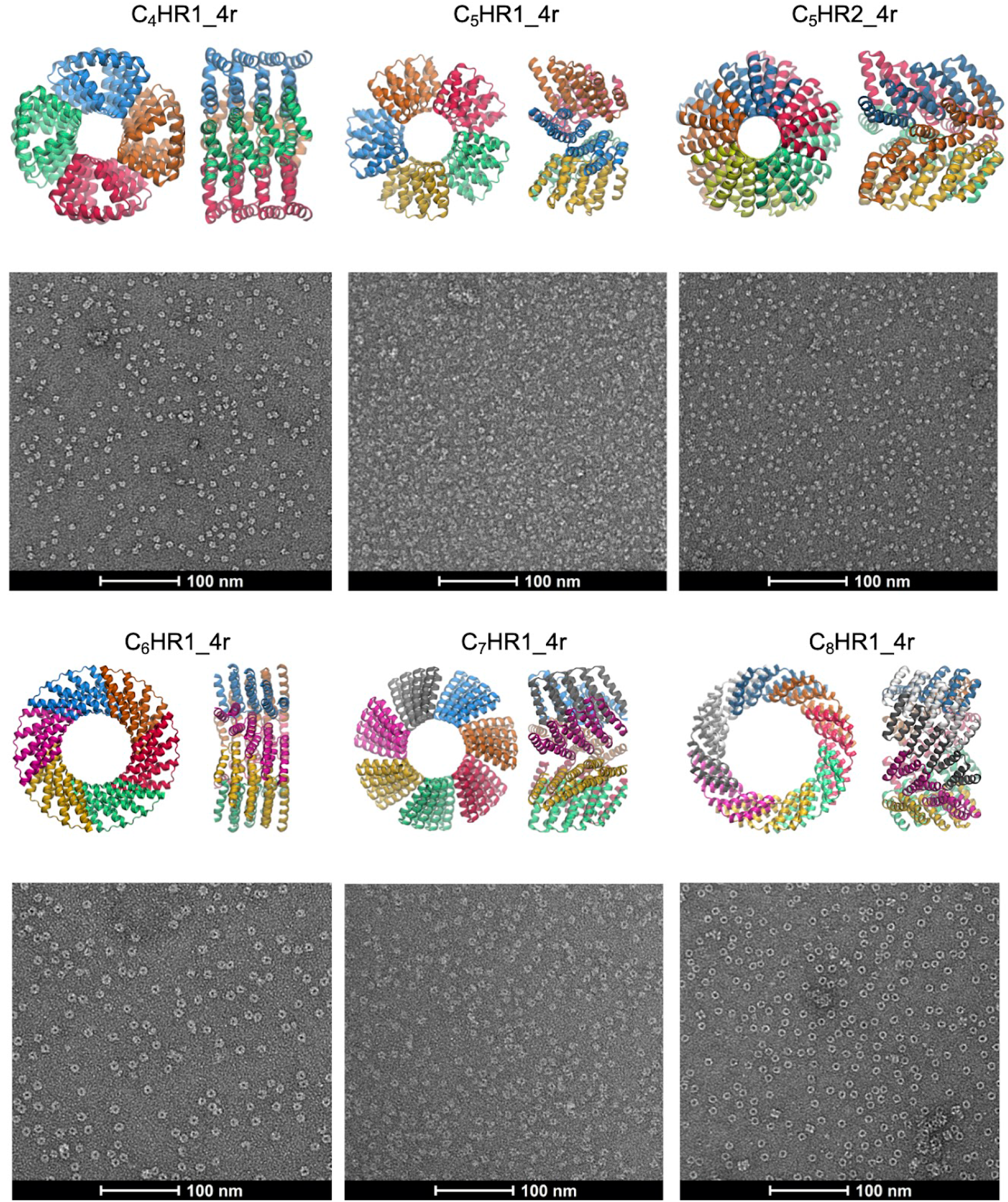
Negative stain electron micrographs of C_4_-C_8_ multiplex assemblies and corresponding design models

**Extended Data Figure 3:**
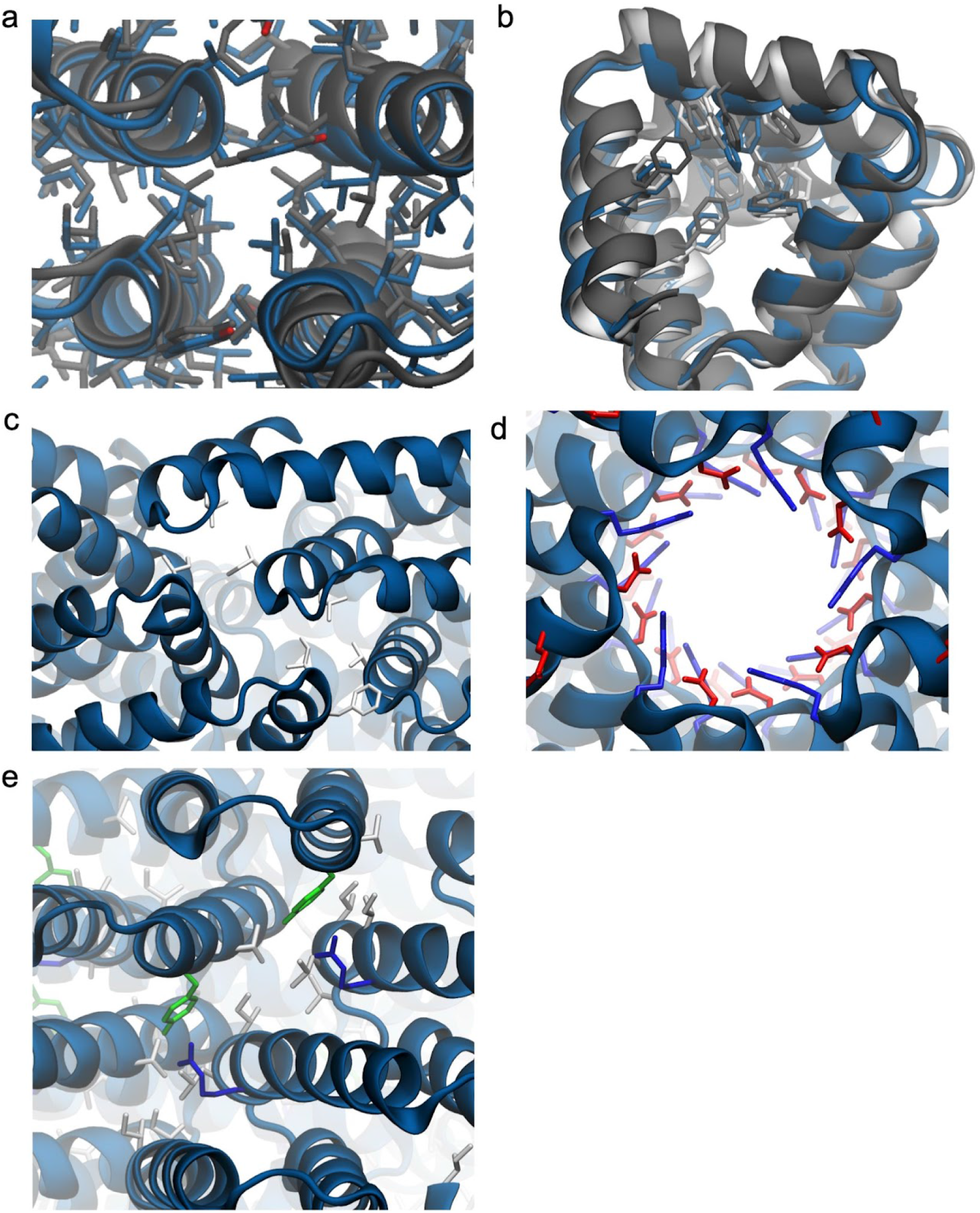
Sidechain rotamers of higher resolution structures. a. C_2_HR1_4r design model (blue) overlaid with crystal structure (gray) zoomed in on the C_2_ interface. The core, interfacial residues for both are shown. b. Buried phenylalanine residues for single chain for C_4_HR1_4r design model (blue), crystal structure (gray) and cryoEM model (white). c. Interfacial residues for C_3_HR3_4r. d. Salt bridging residues at the inner radius of C_5_HR2_4r. Residues are colored blue for basic and red for acidic. e. Interfacial residues for C_6_HR1_4r.

**Extended Data Figure 4:**
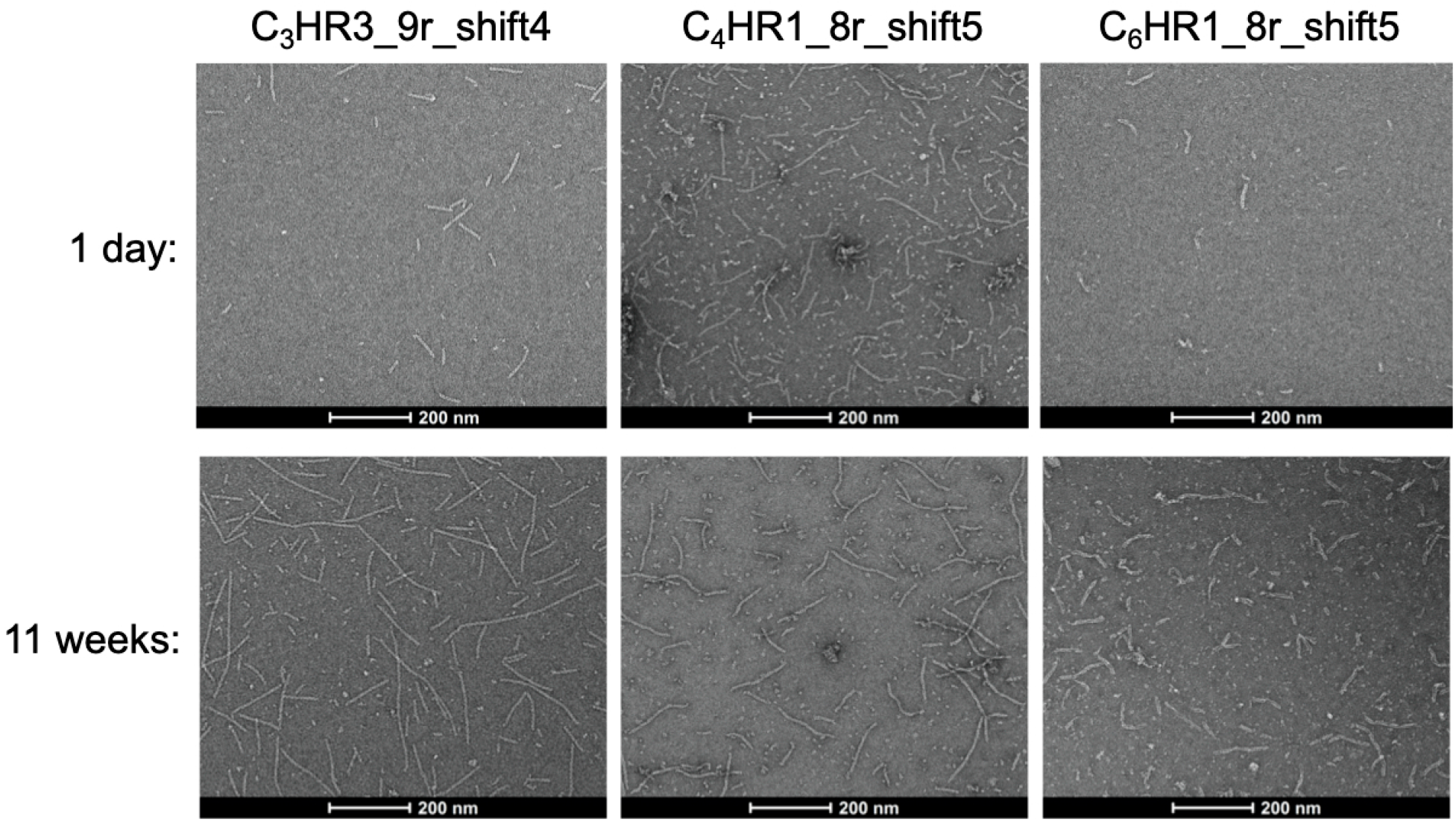
Fiber growth over time at 4°C

**Extended Data Table 1.** SEC-MALS measurements of bounded multiplexes

|  | Expected [kDa] | Observed [kDa] |
| --- | --- | --- |
| **C<sub>2</sub>HR1_4r** | 36.8 | 41.17 (±10.211%) |
| **C<sub>2</sub>HR2_4r** | 46.1 | 40.59 (±20.023%) |
| **C<sub>2</sub>HR3_4r** | 58.7 | 56.94 (±19.229%) |
| **C<sub>3</sub>HR1_4r** | 61.2 | 53.82 (±18.653%) |
| **C<sub>3</sub>HR2_4r** | 67.6 | 68.27 (±21.962%) |
| **C<sub>3</sub>HR3_4r** | 84.7 | 77.47 (±19.192%) |
| **C<sub>4</sub>HR1_4r** | 88.2 | 94.1 (±23.838%) |
| **C<sub>5</sub>HR1_4r** | 99.0 | 93.4 (±20.785%) |
| **C<sub>5</sub>HR2_4r** | 92.5 | 89.71 (±19.229%) |
| **C<sub>6</sub>HR1_4r** | 145.3 | 143.3 ($\pm 14.278\%$) |
| **C<sub>7</sub>HR1_4r** | 160.7 | 146.2 ($\pm 19.392\%$) |
| **C<sub>8</sub>HR1_4r** | 144.2 | 135.2 ($\pm 17.696\%$) |
| **C<sub>3</sub>HR3_8r** | 141.1 | 129.4 ($\pm 18.400\%$) |
| **C<sub>4</sub>HR1_8r** | 171.6 | 168.0 ($\pm 16.541\%$) |
| **C<sub>5</sub>HR2_8r** | 159.5 | 146.9 ($\pm 15.155\%$) |
| **C<sub>6</sub>HR1_8r** | 261.4 | 262.2 ($\pm 14.237\%$) |

**Extended Data Table 2:** Analysis of SAXS scattering curves

|  | **Radius of gyration**<br>**(model) [Å]** | **Radius of**<br>**gyration**<br>**(experiment) [Å]** | **I(0) [a.u.]** | **V<sub>r</sub>** | **$\chi^2$** |
| --- | --- | --- | --- | --- | --- |
| **C<sub>2</sub>HR1_4r** | 19.74 | 26.1 | 6.709 | 6.511 | 0.445 |
| **C<sub>2</sub>HR2_4r** | 22.313 | 28.302 | 29.978 | 7.973 | 1.997 |
| **C<sub>2</sub>HR3_4r** | 22.317 | 25.319 | 30.194 | 3.087 | 0.623 |
| **C<sub>3</sub>HR1_4r** | 22.552 | 25.904 | 106.523 | 4.044 | 2.777 |
| **C<sub>3</sub>HR2_4r** | 28.436 | 24.530 | 23.417 | 6.164 | 0.175 |
| **C<sub>3</sub>HR3_4r** | 27.742 | 29.494 | 102.630 | 3.964 | 1.447 |
| **C<sub>4</sub>HR1_4r** | 26.879 | 32.472 | 78.983 | 4.393 | 1.495 |
| **C<sub>5</sub>HR1_4r** | 28.123 | 31.799 | 32.880 | 6.194 | 0.384 |
| **C<sub>5</sub>HR2_4r** | 26.587 | 32.513 | 32.390 | 12.223 | 0.933 |
| **C<sub>6</sub>HR1_4r** | 35.314 | 41.314 | 45.863 | 6.317 | 1.514 |
| **C<sub>7</sub>HR1_4r** | 33.968 | 37.277 | 60.060 | 5.173 | 0.556 |
| **C<sub>8</sub>HR1_4r** | 39.128 | 55.374 | 52.849 | 8.808 | 7.031 |

**Extended Data Table 3:**
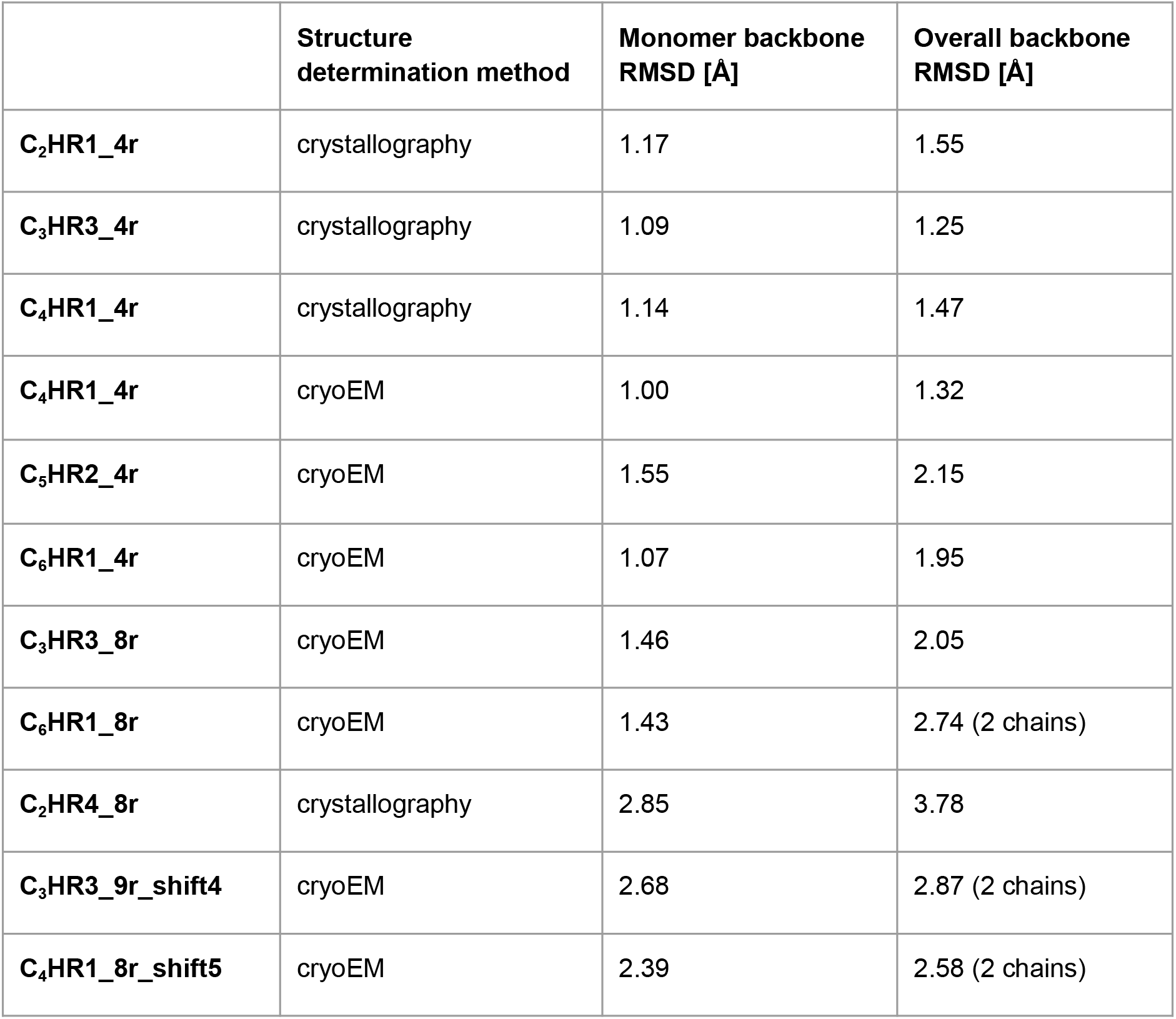
RMSDs for designs models vs high resolution structures

