## Supplementary Materials for "Precisely patterned nanofibers made from extendable protein multiplexes"

### Materials and Methods

#### Computational design of four repeat multiplexes

The design protocol for the repeat protein multiplexes is an adapted and expanded protocol derived from Brunette *et al*. 2015 (3). Repeat protein monomers were generated using the Rosetta RemodelMover. A blueprint file containing the specified secondary structure for a single repeat is provided, and the mover was specified to propagate and link this repeat four total times. Permutations of helix lengths between 9 and 22 and loop lengths between 1 and 4 were attempted for both two and three helix repeats. Additional constraints were placed to ensure helix-helix contacts between neighboring repeats. Following the RemodelMover, the FixAllLoops mover was used to replace any distorted loops that may have been introduced during the remodel step. All backbone monomers were filtered by motif score (3) and the worst9mer filter. For the motif score filter, a threshold of -3.5 was used. For worst9mer, a cutoff of 0.15 for helices and 0.4 overall was used.

Satisfactory backbones were propagated to 12 repeats and aligned so that the helical axis of the monomer aligned with the z-axis. Once aligned, the monomer was copied around the z-axis. Two to eight total copies were attempted, and copy numbers where there were no clashes, but had helix-helix contacts were selected. Sequences were first painted onto the backbones using the Rosetta FastDesign mover. Helical symmetry was enforced to maintain identical sequence and backbone conformation between repeats and other chains. Once the fully symmetric sequence is designed, the system was cut down approximately four repeats per chain. For the two helix repeats, the chain could be cut at either the first or the second loop at the start and the end of the monomer. All four permutations were generated. For the three helix repeats, all nine permutations were generated. A final sequence design of the protein surface was done to remove any hydrophobic patches exposed after backbone truncation. For this step, only cyclic symmetry was applied. Additionally, the surface aggregation potential (SAP) score constraint was used to further minimize hydrophobic patches on the surface. For the Rosetta designed multiplexes, 56 were tested. Of these 56, 27 were soluble, 4 were confirmed to have the correct oligomeric state by SEC-MALs and one ($C_2$HR1_4r) was validated by SAXS (Table S4). In order to increase the success rate of our designed multiplexes, we used machine learning methods to redesign the sequences.

#### Protein design rescue by the machine learning method: ProteinMPNN

Designs output from the method above were redesigned by proteinMPNN (5). Tie constraints, which enforce that pairs of residues are identical, were used such that repeat symmetry of the buried core residues and cyclic symmetry across chains was

maintained. Four sequences for each backbone were generated. All designs except $C_2$HR1_4r and $C_2$HR4_8r were redesigned using proteinMPNN. The code used to generate these sequences can be found here:
https://github.com/nbethel/CHR_multiplexes

### Protein folding validation by Alphafold2 and Alphafold multimer

In order to verify that the designed sequences would fold into the correct structure, each protein structure was predicted by Alphafold multimer. Model 1 was used for all predictions. Alpha carbon RMSD of predictions to the design model was used to select designs for experimental screening. $C_8$ designs were generally too large to be reliably predicted using Alphafold multimer. For these assemblies, we used alphafold2 with three chains. We input the design model as the initial guess, as this helps to find a correct solution for predicting multiple chains with alphafold2. For alphafold2, the default model 4 was used. It is important to note that all models give largely the same answer for these designs, so the choice of model 4 was arbitrary. Designs were again selected according to 2 Å alpha carbon RMSD to the design model.

### Extension of multiplexes from four to eight repeats

A subset of the four repeat designs that were experimentally validated were redesigned as eight repeat versions. The backbone was first propagated to ten repeats, then the chain was cut to approximately 8 repeats. All permutations of different cut points were attempted, as was done for the four repeat multiplexes. For all extensions, the entire sequence was redesigned using proteinMPNN. Internal repeat symmetry and cyclic symmetry was enforced. Additionally the four inner repeats were subjected to full repeat symmetry. This was done to enable the propagation of these assemblies by copying the internal sequence without any computational redesign of the sequence. All designs were evaluated by either Alphafold2 or Alphafold multimer before ordering for experimental characterization.

### Patterned fiber design

To generate patterned fibers, multiplexes were extended to lengths between six and twelve repeats. Adjacent monomers were offset along the helical axis in increments of repeat height and rotation. Using these staggered monomers as reference, helical symmetry was applied to generate copies that extend unbounded along the fiber axis. Once the fiber geometry was established, proteinMPNN was used to generate sequences while maintaining internal repeat symmetry with each monomer and helical symmetry across monomers. Fibers with suitable helix-helix contacts and absent clashes were selected for experimental characterization.

### Preparation of genes from computational designs

Monomers were reverse translated using domesticator (https://github.com/rdkibler/domesticator). These genes were ordered either by IDT or genscript and inserted in pET29b+ vector at NdeI and XhoI restriction sites.

### Buffers and media

LB: 1.2% [wt/vol] tryptone, 2.4% [wt/vol] yeast extract, 0.4% [vol/vol] glycerol, 17 mM KH2 PO4, 72 mM K2 HPO4

TBM-5052: 2.4% [wt/vol] yeast extract, 1.2% [wt/vol] tryptone, 0.5% [wt/vol] glycerol, 0.05% wt/vol D-glucose, 0.2% wt/vol D-lactose, 25 mM Na2HPO4, 25 mM KH2PO4, 50 mM, NH4Cl, 5 mM Na2SO4, 2 mM MgSO4, 10 μM FeCl3, 4 μM CaCl2, 2 μM MnCl2, 2 μM ZnSO4, 400 nM CoCl2, 400 nM NiCl2, 400 nM CuCl2, 400 nM Na2MoO4, 400 nM Na2SeO3, 400 nM H3BO3

Lysis buffer: 25 mM Tris pH 8, 300 mM NaCl, 20 mM Imidazole
Elution buffer: 25 mM Tris pH 8, 300 mM NaCl, 500 mM Imidazole
SEC running buffer: 25 mM Tris pH 8, 300 mM NaCl

### Protein expression and purification

Plasmids were transformed into either lemo21 or bl21de3 expression competent *E.Coli* cells. Transformed colonies were expressed by 50 mL 24hr autoinduction. The cultures were lysed by sonication and purified using Ni-NTA immobilized metal affinity columns. Monodisperse designs that were identified as soluble by SDS polyacrylamide gel electrophoresis were purified further by SEC. For SEC, an akta machine was used with a GE Superdex 200 30x100 GL.

### Characterization by SEC-MALS and SAXS

Multiplexes identified as soluble and monodisperse by SEC were further characterized by SEC-MALS. A volume of 100 uL was injected into an Agilent 1200 HPLC fitted with a Wyatt Heleos DAWN light scattering detector, and a Wyatt Optilab rEX refractive Index detector. A GE Superdex 200 10×300 was used with Pierce 20mM Tris 150mM NaCl pH 8 running buffer, ASTRA 7.0 used for analysis.

SAXS measurements were carried out by the SYBYLIS group. Frameslice was used to preprocess the SAXS scattering data. To get a more realistic matching to experiment, histidine tags were added to the protein structures using Alphafold multimer, model 1. Alphafold multimer, model 1 returned nonphysical, backbone clashing solutions for the $C_5$-$C_8$ multiplexes. Model 3 returned non clashing solutions for $C_5$HR1_4r, $C_5$HR2_4r and $C_7$HR1_4r, so these predicted structures were used in lieu of the model 1 prediction. For $C_6$HR1_4r and $C_8$HR1_4r, no models produced non clashing solutions. For these designs, the single chain predictions were generated using Alphafold2. The generated monomers were copied and aligned to the original design models. The SAXS curves for each HIS tagged model was calculated using the command line implementation of FoXS.

### X-ray crystallography

SEC purified samples were concentrated 15 to 50 mg/ml and crystallization plates were set up using a Mosquito from SPT Labtech, then imaged using UVEX microscopes and UVEX PS-600 from JAN Scientific. Initial trials were carried out using the JCSG I-IV, JCSG+, Morpheus and Classics1-2, and MPD screens, then optimized as needed. For $C_2$HR1_4r crystals grown in 0.1 M MES pH 6.0, 3.2 M Ammonium sulfate and diffraction data was collected at the Berkeley Center for Structural Biology at the Advanced Light Source (ALS). For $C_4$HR1_4r crystals grown in 12.5% *w/v* PEG 1000, 12.5% *w/v* PEG 3350, 12.5% *v/v* MPD, 0.02*M* of carboxylic acids, & 0.1*M* MOPS/HEPES-Na pH7.5 and for $C_3$HR3_4r crystals grown in 0.1 M Imidazole.HCl pH 8.0 15% (w/v) MPD; 5% (w/v) PEG 4000. Diffraction data collected at the Northeastern Collaborative Access Team (NE-CAT) facility at the Advanced Photon Source at Argonne National Laboratory (APS). For $C_2$HR4_8r, crystals were grown using sitting drop vapor diffusion by mixing protein and crystallization solution (0.1M Tris-HCl pH 8.5, 25%(w/v) PEG 3000) in a 1:1 ratio.

X-ray intensities and data reduction were evaluated and integrated using XDS (14) and merged/scaled using Pointless/Aimless in the CCP4 program suite (15). Structure determination and refinement starting phases were obtained by molecular replacement using Phaser (16) using the designed model for the structures. Following molecular replacement, the models were improved using phenix.autobuild (17); efforts were made to reduce model bias by setting rebuild-in-place to false, and using simulated annealing and prime-and-switch phasing. Structures were refined in Phenix (17). Model building was performed using COOT (18). The final model was evaluated using MolProbity (19). Details of data collection and refinement can be found in Table S5. Data deposition, atomic coordinates, and structure factors reported in this paper have been deposited in the Protein Data Bank (PDB), http://www.rcsb.org/ with accession codes $C_2$HR1_4r (8EOV), $C_3$HR3_4r (8EOZ), $C_3$HR1_4r (8EOX) and $C_2$HR4_8r (8ERW).

### Negative stain electron microscopy

SEC purified samples were diluted to ~0.01 mg/ml using SEC buffer immediately before sample application to glow discharged Gilder grids overlaid with a thin layer of carbon (Electron Microscopy Sciences). Grids were then stained using 2% uranyl formate for 2 minutes. Dried grids were screened on a 120 kV Talos L120C transmission electron microscope. The *E. Pluribus Unum* (EPU) (FEI Thermo Scientific) software was used for automated data collection. Two dimensional class averages and three dimensional maps were generated using cryosparc (20)

### CryoEM sample preparation, data collection, and analysis

Protein samples were prepared by diluting or concentrating to 0.5 to 2.0 mg/ml. For the $C_4$HR1_4r, $C_5$HR2_4r and $C_6$HR1_4r, 2.0 uL of sample was applied to glow discharged CF-2/2-4C-T grids (Electron Microscopy Sciences). For $C_4$HR1_8r, $C_5$HR2_8r, $C_6$HR1_8r, $C_3$HR3_9r_shift4 and $C_3$HR1_8r_shift5, 3.0 uL of sample was applied to glow discharged 300 mesh copper quantifoil R 2/2 UT grids (Electron Microscopy Sciences). Using a Vitrobot Mark IV (FEI Thermo Scientific), samples were blotted with either -1 or 0N blot forces from 0.5 to 7.5 s and plunge frozen liquid ethane. All grids were screened and collected on a 200kV Glacios transmission electron microscope (FEI Thermo Scientific) fitted with a Gatan K3 Summit direct electron detector. Movies were collected using the automated software serialEM (21), at 0.05 frames/sec for 99 frames with a dose of 50 e/Å$^2$.

Data processing of the cryoEM micrographs was carried out using cryosparc (20). Movies were motion corrected using “Patch frame motion correction”, and contrast transfer functions were calculated using “Patch CTF estimation”. Images were manually curated to remove images with poor CTF fits and ice quality. For the bounded designs, particles were first selected using “Blob picker”, then resulting class averages were used as templates for the “Template picker”. Class averages were obtained using the “2D class” function. Selected 2D classes were used as input for “3D *ab initio*” reconstructions, then passed “Non uniform refinement” with symmetry applied to obtain the final maps. For the fibers, the “Filament tracer” function was used to pick fibers from the images. The fibers were then class averaged using the “2D class” function, then filament initial reconstructions were generated using the “Helical refinement” tool. Helical parameters were estimated using the “Symmetry search utility”, and these parameters were input for a final round of “Helical refinement” with symmetry applied. Local resolution estimates were determined in CryoSPARC using an FSC threshold of 0.143.

### CryoEM model building and validation

The *de novo* predicted design models for each design (reported here) were used as initial references for building the final cryoEM structures. The models were manually edited and trimmed using Coot (22,23). We further refined each structure in Rosetta using density-guided protocols (24). EM density-guided molecular dynamics simulations were next performed using Interactive Structure Optimization by Local Direct Exploration (ISOLDE) (25), with manual local inspection and guided correction of rotamers and clashes throughout simulated iterations. ISOLDE runs were performed at a simulated 25 Kelvin, with a round of Rosetta density-guided relaxation performed afterward. This process was repeated iteratively until convergence and high agreement with the map was achieved. Multiple rounds of relaxation and minimization were performed on each design, followed by human inspection for errors after each step. Throughout this process, we applied strict non-crystallographic symmetry constraints in Rosetta (26). Phenix real-space refinement was subsequently performed as a final step before the final model quality was analyzed using Molprobity (27) and EM ringer (28). The only deviation to this pipeline was with $C_6$HR1_8r, which deviated significantly from the design model. Monomers for $C_6$HR1_8r were first rigid-body docked individually into the C7 cryoEM map using Chimera (29), followed by an initial round of Rosetta using density-guided protocols. Following this, the model was iterated and finalized similarly to the other six structures. Figures were generated using either UCSF Chimera or UCSF ChimeraX (30).

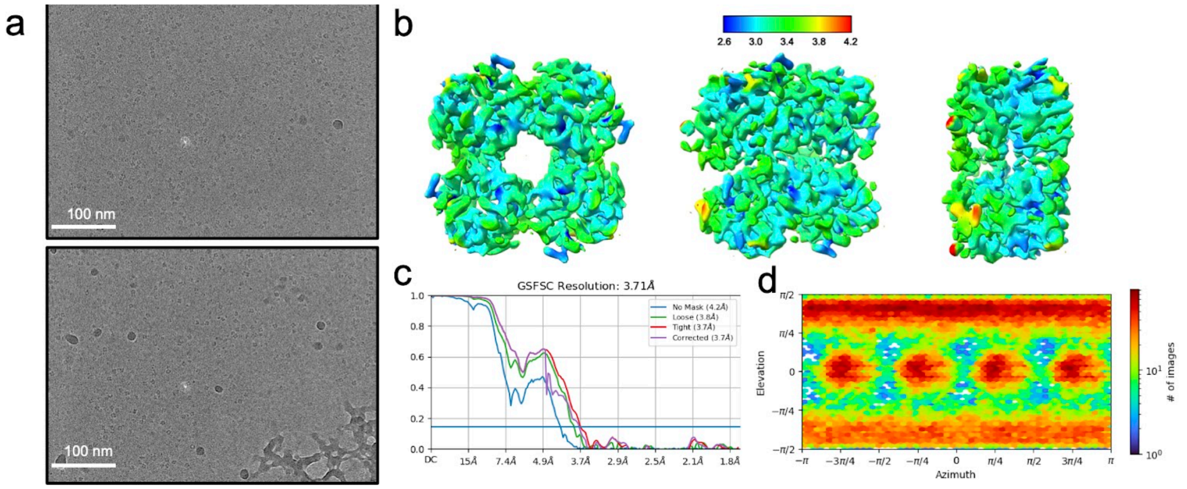


Figure S1: CryoEM micrographs and processing plots for $C_4$HR1_4r. a. Representative raw micrographs. Local resolution overlaid on density isosurface. c. Gold standard Fourier shell correlation curves. d. Viewing direction distribution

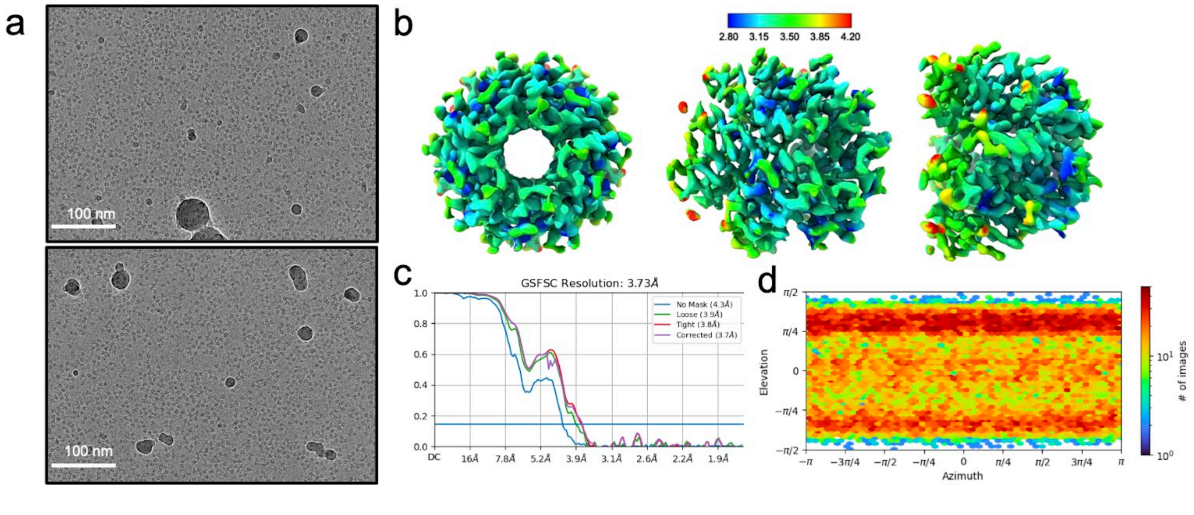

Figure S2: CryoEM micrographs and processing plots for $C_5$HR2_4r. a. Representative raw micrographs. Local resolution overlaid on density isosurface. c. Gold standard Fourier shell correlation curves. d. Viewing direction distribution

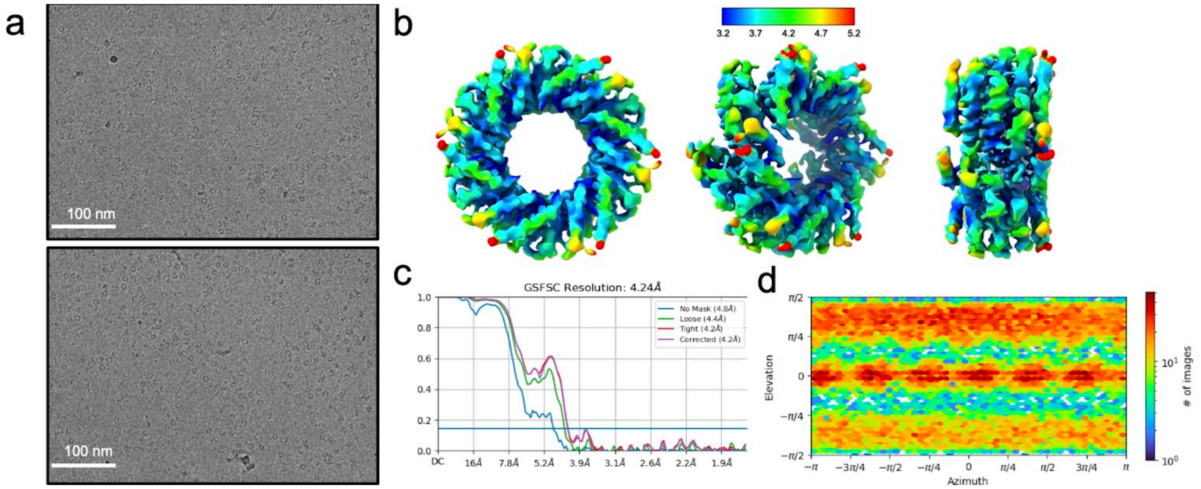

Figure S3: CryoEM micrographs and processing plots for $C_6$HR1_4r. a. Representative raw micrographs. Local resolution overlaid on density isosurface. c. Gold standard Fourier shell correlation curves. d. Viewing direction distribution

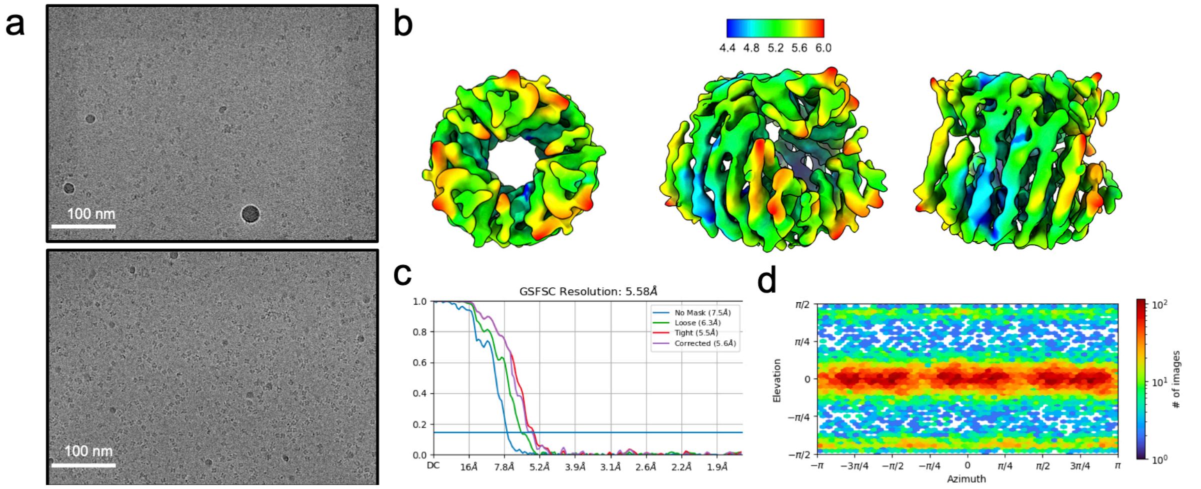

Figure S4: CryoEM micrographs and processing plots for $C_3$HR3_8r. a. Representative raw micrographs. Local resolution overlaid on density isosurface. c. Gold standard Fourier shell correlation curves. d. Viewing direction distribution

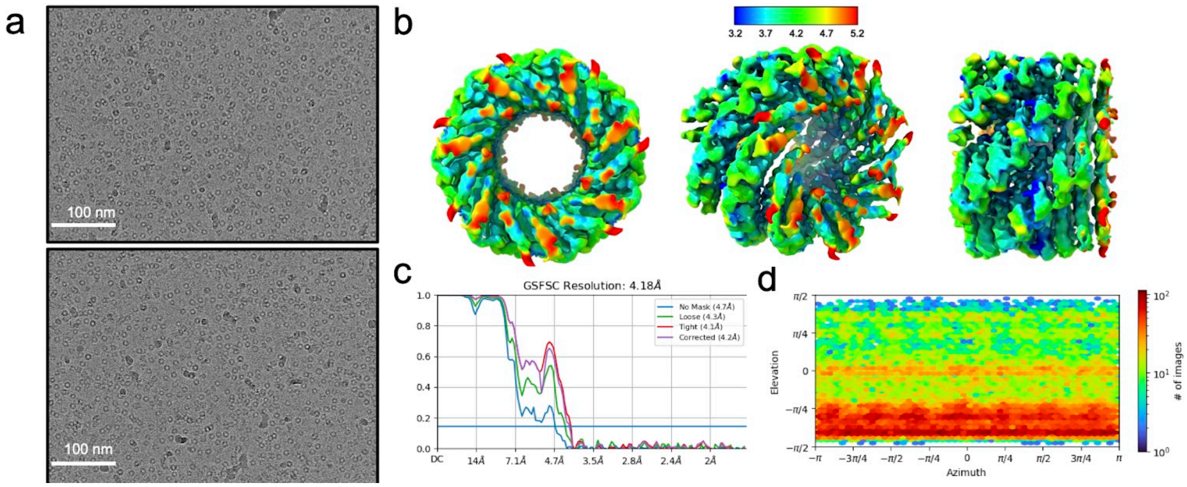


Figure S5: CryoEM micrographs and processing plots for $C_6$HR1_8r. a. Representative raw micrographs. Local resolution overlaid on density isosurface. c. Gold standard Fourier shell correlation curves. d. Viewing direction distribution

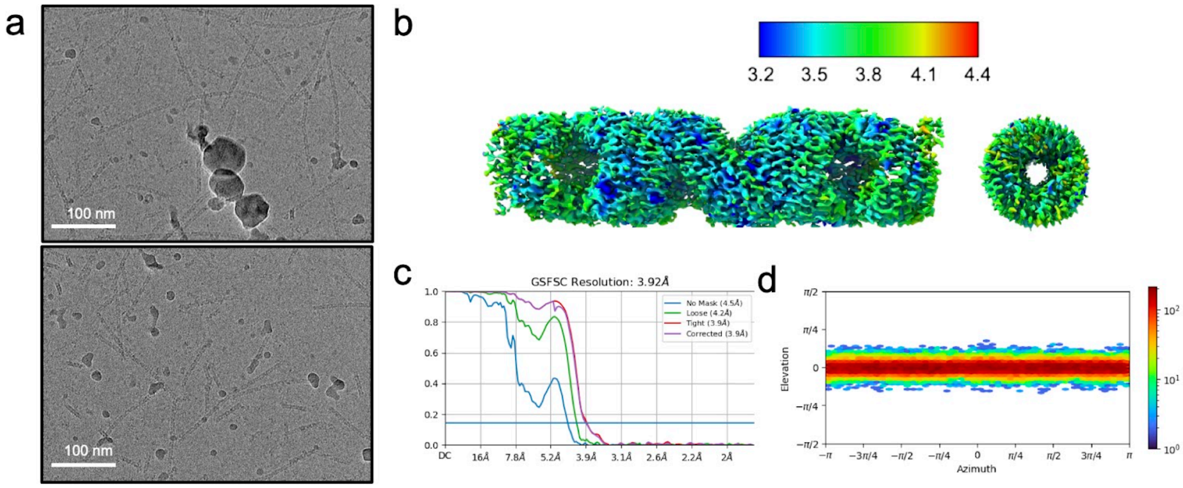


Figure S6: CryoEM micrographs and processing plots for $C_3$HR1_9r_shift4. a. Representative raw micrographs. Local resolution overlaid on density isosurface. c. Gold standard Fourier shell correlation curves. d. Viewing direction distribution

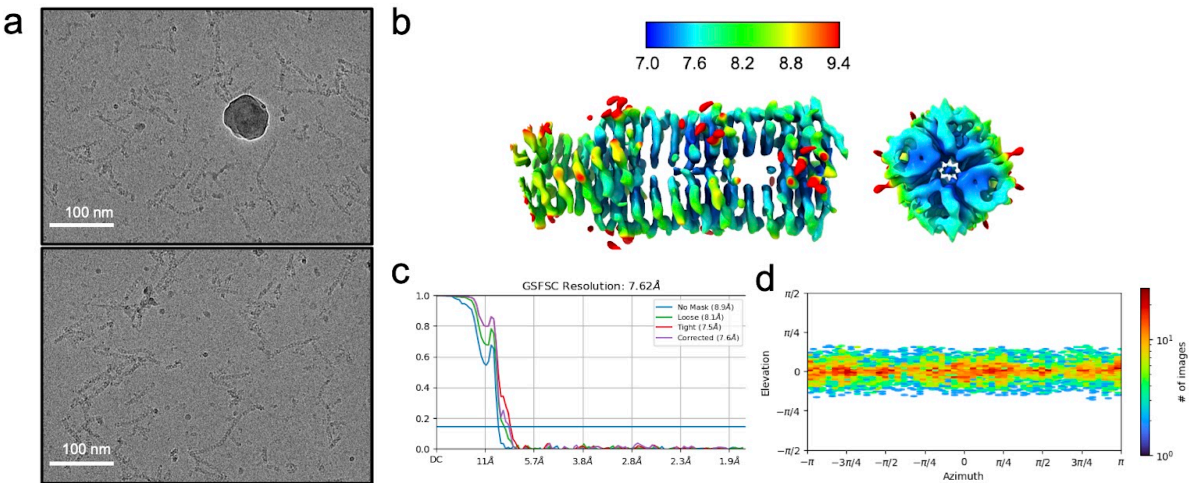


Figure S7: CryoEM micrographs and processing plots for $C_4$HR1_8r_shift5. a. Representative raw micrographs. Local resolution overlaid on density isosurface. c. Gold standard Fourier shell correlation curves. d. Viewing direction distribution

Table S1: Symmetry, sequences, solubility and oligomerization checks for presented CHR multiplexes. A selection of 10 proteinMPNN (mpnn1-10) and 10 rosetta (rosetta1-10) designed sequences for designs the solubility, oligomeric state or SAXS validation are also included, for comparison.

| | **Cyclic symmetry** | **Sequence** | **Soluble** | **Correct oligomeric state** |
|---|---|---|---|---|
| **$C_2$HR1_4r** | 2 | MDYQEALLELIERLLRKLNVDPRDIKRIEQQLRDLDI YQIALLLLIILLLRKLNVDPRDIKRILQQLIDLDIYQIAL LLLIILLLHKLNVDPRDIKRILQQLIDLDIEQIAELLLRIL ELRKRNEDPRDIKRELQQLIDDDLEHHHHHH | yes | yes |
| **$C_2$HR2_4r** | 2 | MMREELLAKLVPESLRWMRELLRETCERLVERGD EVAQELLDVMEALEEDEERFLEHLKLVPLSEQWME MLLFLTCLALVEEGDEVARKLLEVHVELLKDKERFR ELLKLVPSSLRWMEQLLELTCLSLVMEGDEVAREAL EVLIELREDEEFFRELLELVPRGLRWLEILFFLTCLAL MREGDEEARALLRERIELLRDREAFLRYLKEVPLER RTEELLERIRRLRKRMGSWLEHHHHHH | yes | yes |
| **$C_2$HR3_4r** | 2 | MMYEEFFERLRAEDPRLRELERKEEEVREKDPEIA FAYESTKENIDVLIPEEYKKWLLDELFNDPRLRELFV KAKEVEVENPKIAIAYNSTIVNLLVLIPKEYRKWFLDL LKNDPRLRELFVKFIEVAVENPKIAIAYLSTIVNLLVLK PEEELEKELNALFNDPVLREEFVEEIRVAVENPKEAI KKKIEEVIKLLEGSWLEHHHHHH | yes | yes |
| **$C_3$HR1_4r** | 3 | MERVKEVRETMDEFIAVMAGRDPEEIKKVVEKLKE LHKTGTPTTRVIIVDKVMEAFRMVMAGRDPESIKHV VELLKWLHKHGDPRHRVIIVRKTMDTFIDVMAGQD PESIKHVVELLKWLYEHGTPNERVIIRRKVREAAEIV KEGGDPESLAAVEELLAWLEEGSWLEHHHHHH | yes | yes |

| | | | | |
|---|---|---|---|---|
| **$C_3$HR2_4r** | 3 | MELEKLKEMWEKSCSPGMVEAFALLDELLTPEERAAIKRIAENMDEELQKLFLMYLLSCSIGEIYAEILLLKELTPEERKAIFEFIENTDKELMKLLLMYKLSRSIGMLYAYLELLRELTPEERKAIFDLLLNMDKEEMEDFLLKLLSQSIGEIYRELRKLRELSPEERREIFKKKEGSWLEHHHHHH | yes | yes |
| **$C_3$HR3_4r** | 3 | MEEVKKKLEEVWKKAKEDAGDNEKFLELLELILENPEILEILELYVFINKEDVVEKLFDVIKKAVEDAGDNEKFLELLKEMLSNPEIFEILLEYVYIKKEDVVEKLFEVIKQAVEDAGDNPVFLKLLKKMISNPEIFEILLEYVYIGKEEVVKKFFEVIKQAVEDAGNNPIFLKLLEKIILDPERFKKLLEKVEVGEEEEVKAEFKEIKKAVEEAGNDPIKLKELEEKLGSWLEHHHHHH | yes | yes |
| **$C_4$HR1_4r** | 4 | MMKKELYEFYFMPPLKQIEFLEELVNNPEKFKEFFKRLKEEPGAMELFLRNLYLMHPMVQIYFLELLVENPELFKLFFEYLEECPGAMEQFLLNLYLLHPMVQIEFLKLLVENPELFRLFFEYLKRCPGALELFKEIIELLDPIIQKYLKKLLEENPELKALVKEVEEEGSWLEHHHHHH | yes | yes |
| **$C_5$HR1_4r** | 5 | MSAKEIQDILERAEEVVEKGSIKDFLEVLELVKNCEDEEVRNECIKKLAEAVLKMGDIICFLEVLKLVKECENEEVRNECIKILALAVVKMGDILCFLEVLELVKNCPNEEVRNECIRLLAIAVLKMGSKTALAEVKKLVENCPNEEIREECKKILALAEEEGSWLEHHHHHH | yes | yes |
| **$C_5$HR2_4r** | 5 | MVEELKRKLRQAKEDGDEELLERVKNEMLLLAVVDPRVLVEVLNTAKELGDEEMYKKVKGIMLRLAVVDPRVLVLVLELAEALGDEEMKEKVKNIMLLLAVVDPRVLVLVLELAEELGDEEMKKEVEEILDKLAEVDPRVAVLKEVAKKEGSWLEHHHHHH | yes | yes |
| **$C_6$HR1_4r** | 6 | MEEKIKELEEKVEELVKEALEKKDPAVLKKALVCVYEMKKLGMPNEKLIELLKKLVEVLKKLALERVDPAVLDLALVCVYEMKELGMPNEELIKLLKELVEVLRILALINVDPAVLDKALVCVYLMKELGMPNEELIKLLEELVEVLRILALIRVDKRVLDKAEVCIEEMEELGMPEEKIKELREELKFVREILDKLGSWLEHHHHHH | yes | yes |
| **$C_7$HR1_4r** | 7 | MKEVKEKLKKKLEKCKATGDEQDYMDLMKECEKLAKRGCIRGDLETVDTVLEFMLEVCKATGKPEFYQFLMDTCEELARLGCILGNTKTVGLVLLKMLEVCEETGDPEFYRKLMECCEELALLGCRLGNTVTVALVLFFMLEVCRATGDPEFFERLRRTCERLAALGKELGNEKIVALVEFFIEVVDRAGSWLEHHHHHH | yes | yes |
| **$C_8$HR1_4r** | 8 | MEEVLEELKKRLEEAKEKGDYEEIKKYLGSIAIVENNPEVVLKALEERYEIALLTGDFEGVRKYLGSIAIVKNDPEVVLEALETRYLIALMEGDAEEIRKYLRSIAIVKGDKEEEKKALKTLLLVALMEGDEEAVKEYKEKIKEVGSWLEHHHHHH | yes | yes |
| **mpnn1** | 2 | MMRAREEAAVAADPRITRLVEAYRLLRRALGETDEEVEAVAARLEANPIYRALMEAVLADPRILRLRYAREILRRALGETLSQEVEVFERLLKDPLFQALMRAILADPEILRLVYAYEILRLALGETLEEVLECWEELRKDPRFQ | yes | yes |

| | | | | |
|---|---|---|---|---|
| | | AWMRETLADPARLRRVFEELILQLARGETLAEIVARWKEEGSWLEHHHHHH | | |
| **mpnn2** | 2 | MEELKEKLKEAKETNDIELMKEILREILSDSEIIKEVLKEGLLEKILEVALELAEKYNDIELMKLILETILSHSEIIKVVLEKGLLEKILEVALKLAKEKKDIYLMKLILETILSDKRIIEKVLEKGLLEEILEVALELAEENEDPLLIKLILDLINSDERIKKVVEAKGLYEKILKVKEKLEKKYGSWLEHHHHHH | yes | no |
| **mpnn3** | 3 | MVWEKIKKITLKKVKEEGLTEEELKEILKKIKESPVMKLYRGAKYAGDDVTLDFIEETWKECIEEGKTFEELEKRVEEWWNSPVNRLRRGARYAGDDVTLGFIKLTWDKMKEEGKTLEELEEEVRRWQESPVFALLREALRDGDNVRLGIIRLTVYECIVEGKSLEELEARIAEWRESPVFALLDAALAAGDRATLGFIRATLYEAVVRGESLEELERRVAAAAGSWLEHHHHHH | yes | yes |
| **mpnn4** | 3 | MSKSEKVKNTLEEVKEVMKEGKPEKAYELLKELADEISSDLEFVEFVEETDKENISEYVELTLELVEEFIKAGEPEKAARLLELLASLISSSLNFVLFVEYTKKENIEKLVEQTLELVDIFMEEGKPLLAARLLELLASLISTELQFKIFVEYTKKENIQKYVRKTLELVRIFMEEGKPLLAAELLELIESLISTEEQRKLLEEYTLPENLKEKKELKEELEKIFKEGSWLEHHHHHH | yes | no |
| **mpnn5** | 4 | MFYDVMQWIRPASYHAAMAALQAKNPALAALHAMVEEAAKANPFFAVMLHIRPADYWLALAALMAENPELLALHKLVLDAAMKNRRFAVMLFIRPASYWLALYALMAENPELLALFKLVIEKAVSNPEFLRKLYETPSSVWLFKFAEMAENPELLAAYKEEIEALGSWLEHHHHHH | yes | no |
| **mpnn6** | 4 | MKEHLLELFDEIAFSNKARRYWFELYPEIYEYWKKLIKECSIEKGHLLLIFFEIAFSNLERIYWFLLYPEIYKKWLDLIYECSIDKGHLLLIFFEIAFSNLERIFWFLKYPEIVLKAFELIYECSIEKGHELLLFLEESFSNLKKIFKNLKNPEEFLKELEEIAEGSWLEHHHHHH | no | N/A |
| **mpnn7** | 5 | MFHEQLKEDALELGTNEKLKNAALEALDEIEDPEHRLLIIDALELGKTTTLKLAAIKALMEIEDLEHRTLIIDALELGKTTRLRLAAIEALMEIEDLEHRRLIIRALELGTTTTERLAAIKRLMEIEDLEERRAIIKELEGSWLEHHHHHH | no | N/A |
| **mpnn8** | 5 | MLEELLELIGEADLSGEKEAKEEVIRKLKEYVEEKKKEGVDPRDVLRDLIGLADISGLARAKVFVIKKLKEYVKEMREKGVDPWEVLEELIGLADLSGLALAKVFVIEKLEEYVKEMLEEGVDPDEVYARLVELAELSGLALAKVLVRRWEKELKELKEGSWLEHHHHHH | yes | no |
| **mpnn9** | 6 | MSEIDKIIEEYEKKAKEEGVSKKVIGGIIGWMRRILAAGRPEIAEIIREYGIKASKFGVSPTVTGGIIGWMEEILEAGRPEIAERIREYGLEASAFGVSPRVTGGIMDWMIEILEAGRPDLAEKIARLGLRASARGVSPEEVERIMEEMIREEGSWLEHHHHHH | no | N/A |

| | | | | |
|---|---|---|---|---|
| **mpnn10** | 8 | MRKELTDLMLELCRAEDVEVQLACVDRFLEVTENL<br>DDETRLTELEVLLCLAEDERVQLRCVEAFLEVTESL<br>DVEERLAALKRLLRLCEDPRCQRMCVVAFLYVTEE<br>LDDETRLAALEELRELCEDPRIRKVCDVRAAYVER<br>GSWLEHHHHHH | no | N/A |
| **rosetta1** | 2 | MEEEQLEEKRQEIEKALKKNNIEELIELILELIAKIREI<br>AKTEEEQLEILRQLIEKALKKNNIILLILLILILIALILEIA<br>KTEEEQLEILRQLIEKALKKNNIILLILLILILIALILEIAK<br>TEEEQLEILRQLIEKAEKKNNRILLELLRLILEALQREI<br>AKTGSWLEHHHHHH | no | N/A |
| **rosetta2** | 2 | MKEETEKRIREAKEEVEKRIKRTSDEKQQIEEVQQII<br>KELLLEAIAEGNKEILELVIRIAKELVEKLIKRISDEKL<br>QIEVVQQIIKILLLFAIVVGNKEILELVIRIAKELVEKLIK<br>RISDEKLQIEVVQQIIKILLLFAIVVGNKEILELVERIAK<br>ELVEKLQKRISDEKLQEEVEQQIRKIEQLFAEVVGN<br>LEHHHHHH | yes | yes |
| **rosetta3** | 3 | MSDELRELQRENNIEKAWELFRQNNKEEAEKILEEI<br>YQIIRQGSEWLRLRILQEINLIDAAWELFRQNNKEFA<br>KTILEIIRKIIEQGSEWLRLRILEQINLIDAAWQLFREN<br>NKEFAKIILEIIREIIEKGSEWLRLRILKVINLQDAAEEL<br>KRQNNEEFAKIIEEILQQIIEKGSEELREKIKKESGS<br>WLEHHHHHH | no | N/A |
| **rosetta4** | 3 | MDEQRAVWRELEKRREELKKVLQERDIEQVVRVIQ<br>ELLRRFNLDEQRAVWIILEVLRELLKWVLQEYDIEQ<br>VVRVIQTLLRTFNLDEQRAVWIILEVLRELLKWVLQY<br>YDIEQVVRVIQTLLRTFNLDEQRAVWIILEVLRELLK<br>WVLQYYDIEQVVRVIQHLLRSFNLDEQRAKEIIQEV<br>LRELEKWVRQYEDGSWLEHHHHHH | no | N/A |
| **rosetta5** | 4 | MSEELREKIERDGDEDLREIIERAEKYEKRGNDWA<br>VRQIEKLVEKILKLLEQIRRQGDEDLEEIIKRAIKYVK<br>RGNYWAVEEILELVKKILELKLAIEAQGDEDLKEIIQR<br>AIEYVKQGNSEAVREILKLVEKILQLKLAIEAQGDEE<br>LKKEIQKAIEYVKQGNSEAVKRILKEVEKTLREKLKK<br>QASGSWLEHHHHHH | yes | no |
| **rosetta6** | 4 | MDELALRRELRDPEEHEKIREDTSRVLQQLEEAVQ<br>ARDKRALRILLRDPLIHELIREDTSIVLIVLEIAVQARD<br>VEALRILLRDPLIHELIREDTSIVEIVLEIAVQARDVLA<br>LRILLRDPLIHELIREDTSIVHIVLEIARQARDVLALRIL<br>RRDPLIHELEREDTGSWLEHHHHHH | yes | no |
| **rosetta7** | 5 | MDEEVKEVLEKLKDPEEFRRELEEIQKDSAAIIRVIK<br>RWEKIFEVSKEEFVKVFLEVAKTPELFALILEVIRKN<br>SAAIILVIHILENIFKVNKEEFVKVILEVLKTPYLFALLL<br>EVIRKNSAAIHLVIRILENIFKVNKEEFVKVILEVLKTP<br>YLWALLEEVNKKNDQAQKLARHIEENIEEVNKEAFE<br>EVRKKVGSWLEHHHHHH | yes | no |
| **rosetta8** | 5 | MNQEVERLEEELEKQGIDEQQLRRIRKLLQRLAAL<br>GNQEVVRLILELVEEGIDEQQLRRIVKLLLELAALGN<br>QEVVRLILELVQEGIDEQQLRRIVKLLRRLAALGNQ<br>EVVRLILVLVRDGIDEQQLRRSVKKLEEQAAHGNQE | yes | no |

| | | | | |
|---|---|---|---|---|
| | | EVREELVKEISGSWLEHHHHHH | | |
| **rosetta9** | 6 | MDSEKEHSRLRKDRERTSEQEIKEVLERILWEAVAERDSELLHSILRLIREITSEQEIKEVLERILQLAVALRDSELLHSILRLIREITSEQEIKEVLERILEDAVALRDSELLYSILILIWLITSEQEIKEVLERIREQAEALRDSELEHSIKILKKLITSGSWLEHHHHHH | no | N/A |
| **rosetta10** | 8 | MNQEAEIRKARETLRRIQKTWERGNQEEAIERLLELLIRLIVGGNQEALIEVAEILLELIQKIWERGNQEEAIEILLELLIVLIVGGNQEALIRVAQILLELIQKIWERGNQEEAIEILLELLEVLSVGGNQEALIIVALILLLLIQKIWERGNQEEAHEILEELEEVLREGGNQQAQRIVALIQQLLEQKIRERGNQGSWLEHHHHHH | no | N/A |

Table S2: X-ray crystallography data collection and refinement statistics

| | **$C_2$HR1_4r (8EOV)** | **$C_3$HR3_4r (8EOZ)** | **$C_4$HR1_4r (8EOX)** | **$C_2$HR4_8r (8ERW)** |
|---|---|---|---|---|
| **Resolution range** | 42.56 - 1.59 (1.64 -1.59) | 46.35 - 3.0 (3.10 - 3.0) | 53.79 - 3.3 (3.41 - 3.3) | 37.79 - 2.882 (2.985 - 2.882) |
| **Space group** | P 32 2 1 | R 3 :H | P 1 21 1 | P 21 21 21 |
| **Unit cell** | 49.14 49.14 111.21 90 90 120 | 68.22 68.22 139.03 90 90 120 | 62.75 80.36 66.70 90 112.75 90 | 55.318 55.38 145.134 90 90 90 |
| **Total reflections** | 43250 (4240) | 6210 (328) | 17615 (1797) | 45175 (3066) |
| **Unique reflections** | 21628 (2120) | 4528 (328) | 9068 (907) | 10204 (876) |
| **Multiplicity** | 2.0 (2.0) | 1.4(1.0) | 1.9 (2.0) | 4.4 (3.5) |
| **Completeness (%)** | 99.76 (99.67) | 98.49 (99.90) | 97.04 (98.69) | 95.62 (86.48) |
| **Mean I/sigma(I)** | 23.39 (0.68) | 10.95 (4.09) | 7.22 (0.90) | 7.87 (1.39) |
| **Wilson B-factor** | 31.39 | 31.14 | 135.90 | 84.54 |
| **R-merge** | 0.01344 (1.038) | 0.04045 (0.689) | 0.03538 (0.7009) | 0.1155 (0.6324) |
| **R-meas** | 0.01901 (1.468) | 0.05721 (0.746) | 0.05003 (0.9912) | 0.1295 (0.7334) |

| | | | | |
|---|---|---|---|---|
| **R-pim** | 0.01344 (1.038) | 0.04045 (0.282) | 0.03538 (0.7009) | 0.05655 (0.3591) |
| **CC1/2** | 1 (0.389) | 0.990 (0.851) | 0.997 (0.655) | 0.994 (0.677) |
| **CC*** | 1 (0.749) | 0.998 (0.959) | 0.999 (0.889) | 0.999 (0.898) |
| **Reflections used in refinement** | 21592 (2113) | 4527 (328) | 9035 (903) | 10135 (876) |
| **Reflections used for R-free** | 1995 (199) | 459 (34) | 450 (41) | 1028 (90) |
| **R-work** | 0.2023 (0.3946) | 0.2343 (0.2917) | 0.2947 (0.4327) | 0.2611 (0.3263) |
| **R-free** | 0.2244 (0.3745) | 0.2902 (0.3255) | 0.3516 (0.3932) | 0.2862 (0.3642) |
| **Number of non-hydrogen atoms** | 1254 | 1874 | 5619 | 3157 |

Table S3. CryoEM Data Collection Statistics for Bounded Designs

| | $C_5$HR2_4r | $C_3$HR3_8r | $C_4$HR1_4r | $C_6$HR1_4r | $C_6$HR1_8r |
|---|---|---|---|---|---|
| Microscope | Glacios | Glacios | Glacios | Glacios | Glacios |
| Voltage (kV) | 200 | 200 | 200 | 200 | 200 |
| Detector | Gatan K3 Summit | Gatan K3 Summit | Gatan K3 Summit | Gatan K3 Summit | Gatan K3 Summit |
| Recording mode | Counting | Counting | Counting | Counting | Counting |
| Magnification | 45,000x | 45,000x | 45,000x | 45,000x | 45,000x |
| Movie micrograph pixel size (Å) | 0.4425 | 0.4425? | 0.4425 | 0.4425 | 0.4425 |
| Dose rate ($e^-$/Å$^2$/s) | 10 | 10 | 10 | 10 | 10 |
| No. of frames per movie micrograph | 99 | 99 | 99 | 99 | 99 |
| Frame exposure time (ms) | 0.0505 | 0.0505 | 0.0505 | 0.0505 | 0.0505 |
| Movie micrograph exposure time (s) | 5.0 | 5.0 | 5.0 | 5.0 | 5.0 |
| Total dose ($e^-$/Å$^2$) | 50 | 50 | 50 | 50 | 50 |
| Under focus range (μm) | 1.0 - 2.0 | 1.0 - 2.0 | 1.0 - 2.0 | 1.0 - 2.0 | 0.7-1.8 |
| Number of movie micrographs | 306 | 1810 | 1146 | 694 | 1468 |

Table S4. CryoEM Data Collection Statistics for Unbounded Designs

| | $C_3$HR3_9r_shift 4 | $C_4$HR1_8r_shift5 |
|---|---|---|
| Microscope | Glacios | Glacios |
| Voltage (kV) | 200 | 200 |
| Detector | Gatan K3 Summit | Gatan K3 Summit |
| Recording mode | Counting | Counting |
| Magnification | 45,000x | 45,000x |
| Movie micrograph pixel size (Å) | 0.4425 | 0.4425 |
| Dose rate ($e^-/Å^2/s$) | 10 | 10 |
| No. of frames per movie micrograph | 99 | 99 |
| Frame exposure time (ms) | 0.0505 | 0.0505 |
| Movie micrograph exposure time (s) | 5.0 | 5.0 |
| Total dose ($e^-/Å^2$) | 50 | 50 |
| Under focus range (μm) | 0.7 - 1.8 | 0.7 - 1.8 |
| Number of movie micrographs | 916 | 656 |
